# Phosphatidylcholine Metabolism Controls Alveolar Progenitor Renewal and Pulmonary Fibrosis

**DOI:** 10.1101/2025.09.26.678894

**Authors:** Anas Rabata, Yujie Qiao, Weini Li, Haohua Huang, Shaopeng Yu, Xuexi Zhang, Xue Liu, Ningshan Liu, Guanling Huang, Peipei Liu, Ankita Burman, Ales Hampl, Tanyalak Parimon, Changfu Yao, Cory Hogaboam, Peter Chen, Barry Stripp, Anjaparavanda P. Naren, Robert Damoiseaux, Paul W. Noble, Jiurong Liang, Dianhua Jiang

## Abstract

Idiopathic pulmonary fibrosis (IPF) is a progressive, fatal lung disease marked by alveolar type 2 (AT2) stem cell dysfunction and excessive matrix deposition, with no effective treatments. Recent advances have recognized that AT2 cells act as stem cells, in addition to their role in the production of pulmonary surfactants in the distal alveolar space. We and others have reported a failure of AT2 regeneration and a loss of AT2 cells in IPF. We recently further reported that there is a defect in lipid metabolism in IPF AT2 cells and we discovered a selective loss of lysophosphatidylcholine acyltransferase 1 (LPCAT1) in AT2 cells from IPF, as well as in AT2 cells from bleomycin-injured mice. Pharmacological and genetic experiments confirm that LPCAT1 is required for AT2 cell renewal in 3D organoid assays. AT2 cell-specific *Lpcat1* deletion resulted in reduced AT2 renewal, spontaneous lung fibrosis, and heightened susceptibility to bleomycin-induced fibrosis in mice in vivo. Expression-based high-content drug screening with an LPCAT1 knock-in cell line identified several drug families that upregulated LPCAT1 expression. We further confirmed that anti-malarial artesunate and PLA2 inhibitor ONO-RS-082 increased LPCAT1 mRNA expression, promoted AT2 renewal, and attenuated bleomycin-induced lung fibrosis in mice in vivo. Our findings establish LPCAT1 as a critical regulator of AT2 renewal and lipid metabolism in IPF, suggesting that reactivation of LPCAT1 could offer a novel therapeutic strategy for restoring alveolar progenitor function and mitigating lung fibrosis.

## Introduction

Idiopathic pulmonary fibrosis (IPF) is the most severe form of interstitial lung disease (ILD), with patients typically succumbing within three to five years after diagnosis (*1–4*). Despite extensive efforts to investigate IPF pathogenesis, no effective treatments have been conclusively identified (*3*). Key features of IPF development include continuous epithelial damage encompassing loss of both functional integrity and regenerative capacity (*5–8*). Damaged epithelial cells drive fibroblast activation through epithelial-mesenchymal crosstalk, leading to excessive extracellular matrix production and lung fibrosis (*9, 10*). However, the underlying pathogenesis of IPF is still not fully elucidated.

Alveolar type 2 cells (AT2s) are characterized as stem cells with progenitor function and renewal capacity (*11–13*). Our laboratory and others have reported that, in IPF, AT2 cell numbers decline in the lung, while the remaining AT2s exhibit impaired renewal capacity (*5, 6, 14*). Changes in the structure and surfactant composition of AT2s have been reported in IPF (*15, 16*). However, the molecular mechanisms that sustain AT2 stem cell renewal and protect them from the pathological changes leading to lung fibrosis remain unknown. The main structural components of AT2 cell membranes are phospholipids, which regulate cell shape, transport, and signaling (*17, 18*) and play roles in lung diseases, such as IPF, cystic fibrosis, asthma, and chronic obstructive pulmonary disease (COPD) (*19–21*). Phosphatidylcholine (PC) is the major phospholipid in AT2 cell membranes and in pulmonary surfactant, a lipoprotein complex essential for respiration (*22–24*).

In the lung, PC is synthesized via the de novo Kennedy pathway and recycled through the Lands’ cycle. The latter maintains a dynamic balance between lysophosphatidylcholine (LPC) and PC through de-acylation by phospholipase A2s (PLA2) and re-acylation by lysophosphatidylcholine acyltransferases (LPCATs) (*22, 25–28*). Four LPCAT family members (LPCAT1-LPCAT4) have been identified, each with distinct acyl-CoA selectivities and tissue distribution patterns (*28–31*). LPCAT1 (also known as LPLAT8 or previously AGPAT9) is selectively expressed in the lung (*23, 32–36*). Deletion of *Lpcat1* in mice results in respiratory failure and perinatal death (*23*).

We discovered that LPCAT1 expression is significantly reduced in IPF AT2 cells. This prompted us to investigate its role in AT2 stem cell renewal, the mechanisms by which it regulates AT2 renewal and lung fibrosis, and strategies to therapeutically manipulate LPCAT1 expression. In this study, we determined that LPCAT1 is a critical enzyme in PC remodeling, essential for maintaining mitochondrial function, AT2 progenitor renewal, and the recovery of injured AT2 cells. We further identified distinct compounds that upregulate LPCAT1 expression, restore the function of injured AT2 cells, and attenuate bleomycin-induced lung fibrosis. Collectively, these findings indicate that LPCAT1 represents a potential therapeutic target for progressive fibrotic lung diseases such as IPF.

## Results

### Downregulation of LPCAT1 in IPF AT2s

Our previous work showed lipid metabolism, especially PC metabolism, was dysregulated in IPF AT2 progenitor cells (*19*). Analysis of in-house scRNA-Seq data from freshly isolated epithelial cells (CD31^−^CD45^−^EpCAM^+^) of IPF and healthy lung donors (*5, 19, 37*) showed that LPCAT1 was predominately expressed in AT2 and AT1 cells (Figure 1A), while other LPCAT family members are expressed at much lower levels in AT2 cells (Figure S1).

**Figure 1:**
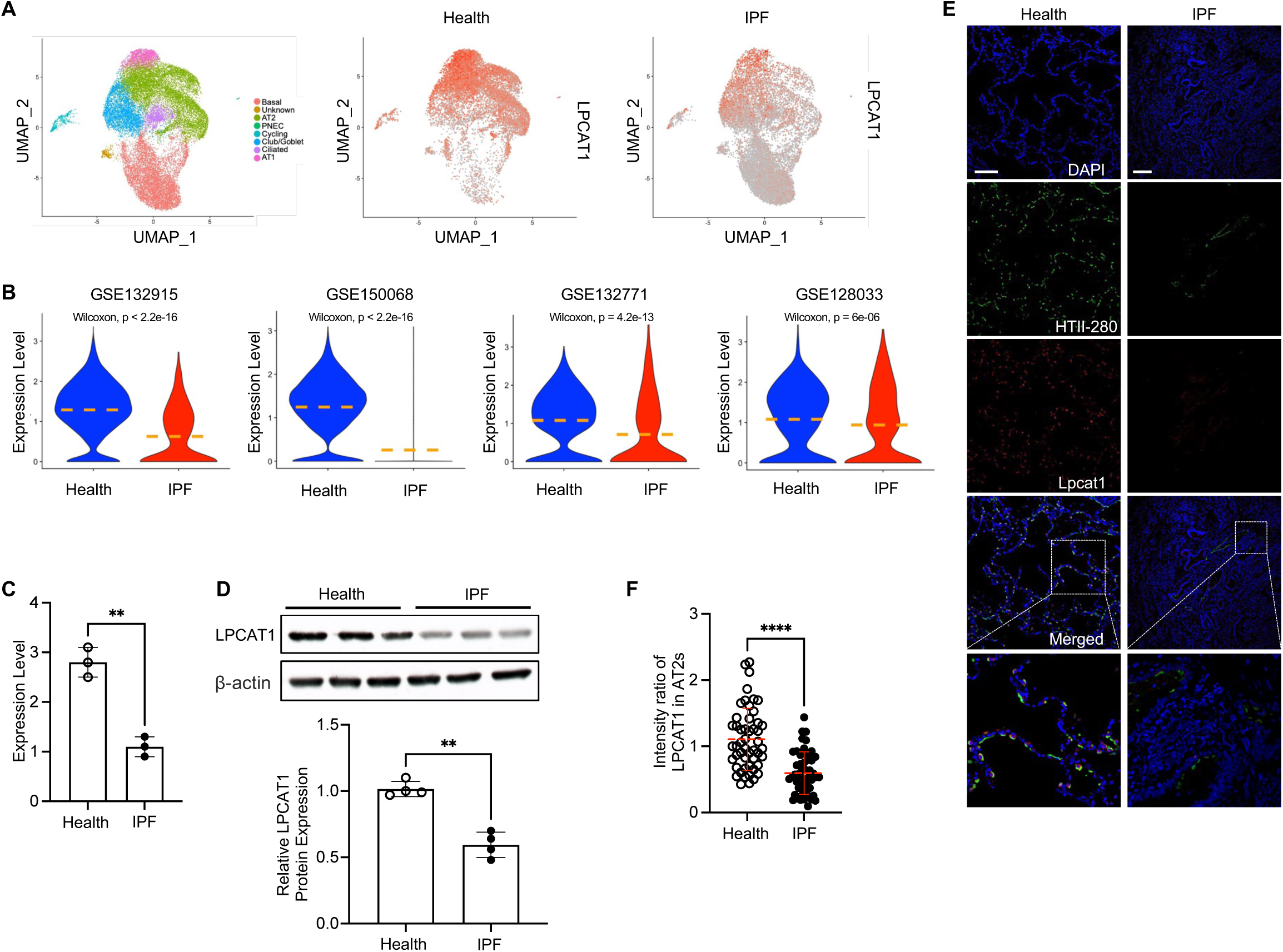
LPCAT1 is predominantly expressed in alveolar epithelial cells and is significantly reduced in IPF. (A) UMAP plots of flow-enriched EpCAM^+^CD31^−^CD45^−^ cells from healthy (11,381 cells, n = 6) and IPF lungs (14,687 cells, n = 6), showing clusters of epithelial cell types and the expression of the *LPCAT1* gene in EpCAM^+^CD31^−^CD45^−^ cells from healthy and IPF lungs from an in-house scRNA-seq dataset. (B) Violin plots showing *LPCAT1* expression levels in AT2s from healthy and IPF lungs from published datasets GSE132915, GSE150068, GSE132771, and GSE128033. (C) Results from qPCR analysis of *LPCAT1* expression in AT2s from healthy and IPF lungs (n = 3 per group, **p < 0.01). (D) Representative photographs and quantification of Western blot analysis of LPCAT1 in healthy and IPF lung tissues, with β-actin serving as a loading control (n = 4 per group, **p < 0.01). (E) Immunofluorescence staining of healthy and IPF lung sections for the AT2 marker HTII-280 (green) and LPCAT1 (red). Scale bars, 100 μm. Staining was performed on lung sections from 4 IPF patients and 4 healthy donors. Representative areas (squares) are shown at higher magnification. (F) Quantification of LPCAT1 staining (relative signal intensity) in individual AT2s (HTII-280+) (50 cells/section, n = 3 per group, ****p < 0.0001).

LPCAT1 expression was significantly reduced in IPF AT2s compared with healthy AT2s (Figure 1A). We confirmed this finding by analyzing additional published scRNA-Seq datasets from IPF patients. LPCAT1 was significantly reduced in IPF AT2s compared with healthy AT2s in GEO datasets GSE132915 (*38*), GSE150068 (*39*), GSE132771 (*40*), and GSE128033 (*41*) (Figure 1B).

To validate LPCAT1 expression in IPF AT2s, we performed qPCR on freshly isolated AT2s (EpCAM^+^HTII-280^+^) from IPF patients and healthy donors (Figure 1C). We further analyzed LPCAT1 protein levels and confirmed significantly lower expression in IPF lung tissues (Figure 1D). Immunofluorescence staining showed that AT2s in IPF lung sections expressed much lower levels of LPCAT1 than those from healthy donors (Figure 1E), accompanied by significantly reduced fluorescence intensity of LPCAT1 staining in AT2 cells from IPF lung sections (Figure 1F).

Next, we investigated *Lpcat* family genes in mouse epithelial cells using in-house scRNA-seq dataset from uninjured and bleomycin-injured mice at multiple time points (*42*). Consistent with human cell analysis, Lpcat1 gene is highly expressed in the AT2 cell cluster (Figure 2A). We analyzed *Lpcat* gene expression in AT2s at Day 4 (the peak of AT2 injury) after bleomycin treatment and compared it with uninjured (Day 0) mice. Lpcat1 expression was significantly downregulated in AT2s from Day 4 bleomycin-injured mouse lungs (Figure 2B). These results were confirmed by qPCR using freshly isolated AT2s (EpCAM^+^CD24^-^Sca-1^-^) from uninjured and bleomycin-injured mice (Figure 2C). Additionally, immunofluorescence staining showed reduced LPCAT1 expression in AT2s from injured mice (Figure 2D), as indicated by lower fluorescence intensity (Figure 2E).

**Figure 2:**
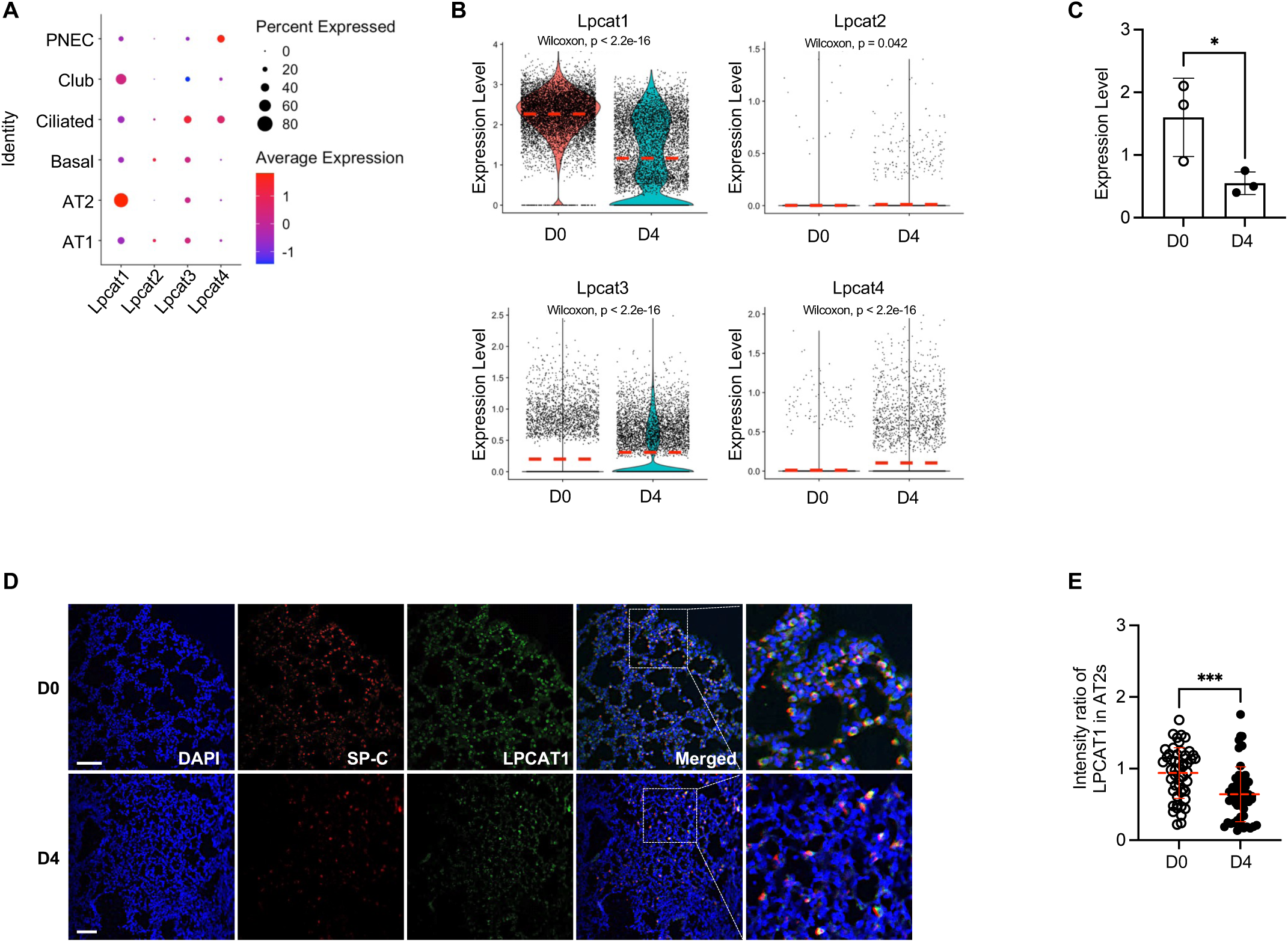
LPCAT1 is predominantly expressed in alveolar epithelial cells cells and is significantly reduced in bleomycin mouse model. (A) Dot plot showing the average expression of *Lpcat* family genes in different clusters of mouse epithelial cell types from an in-house scRNA-seq dataset. (B) Violin plots showing the expression of *Lpcat* family genes in AT2s from mouse lungs of uninjured (D0) (n = 3) and 4 days after bleomycin-injury (D4) (n = 4). (C) Results from qPCR analysis of *Lpcat1* expression in mouse AT2s from uninjured (D0) (n = 3) and 4 days after bleomycin-injury (D4) (n = 3 per group, *p < 0.05). (D) Immunofluorescence staining of mouse lung sections from uninjured (D0) (n = 5) and 4 days after bleomycin-injury (D4) (n = 5). HTII-280 (green), and LPCAT1 (red). Scale bars, 100 μm. Representative areas (squares) are shown at higher magnification. (E) Quantification of LPCAT1 staining (relative signal intensity) in individual AT2s (HTII-280+ cells) (50 cells/section, n = 3 mice per group, ***p < 0.001).

### LPCAT1 is required for AT2 renewal

These expression data suggest that LPCAT1 may play a functional role in maintaining AT2 progenitor cell renewal. We next inhibited LPCAT1 pharmacologically in a 3D organoid culture model. Although no specific LPCAT1 inhibitor exists, the small-molecule compound TSI-01 exerts inhibitory effects on both LPCAT2 and LPCAT1 (*43*). We cultured AT2s with TSI-01 at concentrations ranging from 1 to 20 μM. 3D organoid cultures revealed that TSI-01 significantly inhibited AT2 colony formation efficiency (CFE) in a dose-dependent manner in both mouse and human cells (Figure 3A). This effect was not attributable to reduced cell growth, since 3 and 10 μM TSI-01 showed minimal impact in the AT2 proliferation assay (Figure 3B).

**Figure 3:**
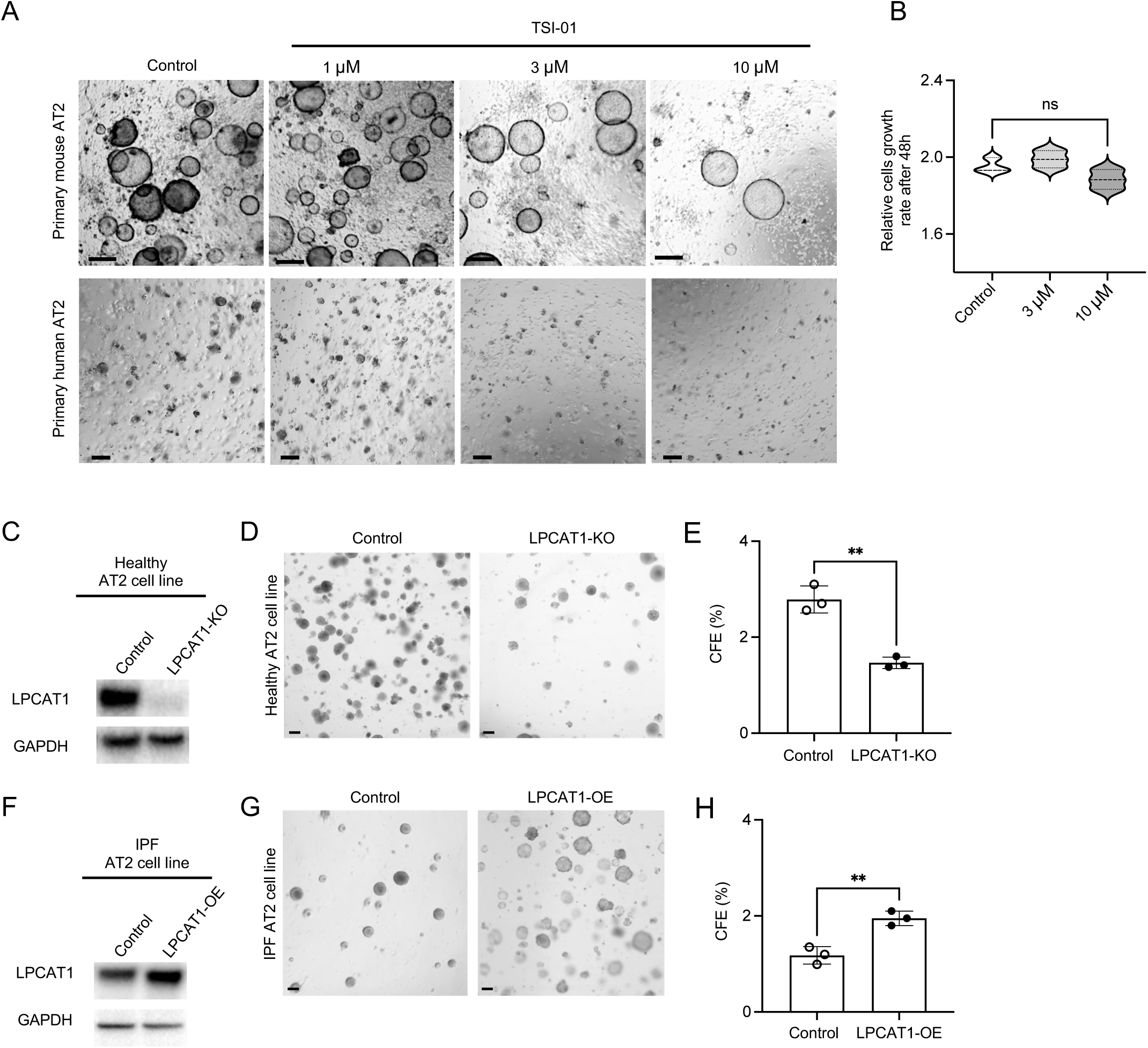
Inhibition of LPCAT1 reduces AT2 renewal. (A) Representative images of 3D organoid cultures (day 14) of primary AT2s from wild-type mice or healthy human lungs in the presence or absence of the Lpcat1 inhibitor (TSI-01). Scale bars, 100 μm. (n = 3 per group). (B) Relative cell growth rate after 48 hours of culture of AT2s in the presence of TSI-01 (3 μM, 10 μM). (n = 3 per group, ns, no significant difference). (C) Representative photographs of Western blot analysis of LPCAT1 after CRISPR-mediated knockout (KO) of *LPCAT1* in AT2 cell lines from healthy lungs (AT2-*LPCAT1*-KO). GAPDH as an internal control, (D) Representative images of 3D organoid cultures (day 14) of AT2-*LPCAT1*-KO cell lines and controls. (E) CFE of AT2-*LPCAT1*-KO cell lines and controls in 3D organoid cultures. (n = 3 per group, **p < 0.01). (F) Representative photographs of Western blot analysis of LPCAT1 after CRISPR-mediated overexpression (OE) of *LPCAT1* in AT2 cell lines from IPF lungs (AT2-*LPCAT1*-OE). GAPDH as an internal control, (G) Representative images of 3D organoid cultures (day 14) of AT2-*LPCAT1*-OE cell lines and controls. (H) CFE of AT2-*LPCAT1*-OE cell lines and controls in 3D organoid cultures. (n = 3 per group, **p < 0.01).

### Genetic gain- and loss-of-function analysis determines the role of LPCAT1 in AT2 renewal

To overcome the limitation that pure primary human AT2s are difficult to maintain and passage, and lack of the ability for genetic manipulation, we adapted a published protocol by Borok and Offringa’s group (*44*) to successfully immortalize several lines of healthy and IPF AT2s. In brief, EpCAM^+^ cells were isolated from both IPF and healthy donor lungs by Magnetic-Activated Cell Sorting (MACS) and subsequently cultured in medium containing the Rho-associated kinase (ROCK) inhibitor. This was followed by lentiviral transduction with SV40 Large T antigen (SV40 LgT), after which HTII-280^+^ cells – a well-established AT2 marker – were selected and sorted (Figure S2).

Western blot analysis showed that LPCAT1 expression was reduced in the AT2 cell lines derived from IPF lungs (Figure S2C). Furthermore, the application of 3D culture conditions resulted in a significantly reduced CFE in AT2s derived from IPF lungs compared to those from healthy lungs (Figure S2D). These findings corroborate our previous observations that LPCAT1 expression is downregulated in AT2s from IPF patients and that these cells exhibit impaired renewal capacity.

To explore the role of LPCAT1 in AT2 cell function, we conducted genetic gain- and loss-of-function (GOF and LOF) analyses using CRISPR/Cas9. In healthy AT2s, LPCAT1 was knocked out (AT2-LPCAT1-KO; Figure 3C), whereas in IPF AT2s, CRISPR activation was employed to enhance LPCAT1 expression (AT2-LPCAT1-OE; Figure 3F). 3D organoid culture models showed that LPCAT1 deletion significantly reduced the CFE of healthy AT2 cell lines (Figure 3D and 3E), which is consistent with our in vivo findings in Lpcat1^AT2^ mice. Conversely, overexpression of LPCAT1 in IPF AT2 cell lines led to a significant enhancement of CFE (Figure 3G and 3H). Our findings identify LPCAT1 as a critical determinant of AT2 renewal.

### LPCAT1 is required for AT2 maintenance in mice in vivo

To elucidate the role of LPCAT1 in AT2 renewal and lung fibrosis in vivo, we generated a *Lpcat1*^flox/flox^ mouse line by introducing loxP sites flanking exon 3 via CRISPR/Cas9 system (Figure 4A), which was crossed it with *Sftpc*-CreER mice (*12*). This cross generates *Sftpc*-CreER;*Lpcat1*^flox/flox^ mice (hereafter referred to as *Lpcat1*^AT2^), in which tamoxifen treatment deletes LPCAT1 in AT2s. The genotype of *Lpcat1*^AT2^ mice was validated using PCR. Histological and immunofluorescence studies revealed that tamoxifen treatment led to the loss of LPCAT1 expression in AT2s, without inducing overt inflammation or histological abnormalities (Figure 4B).

**Figure 4:**
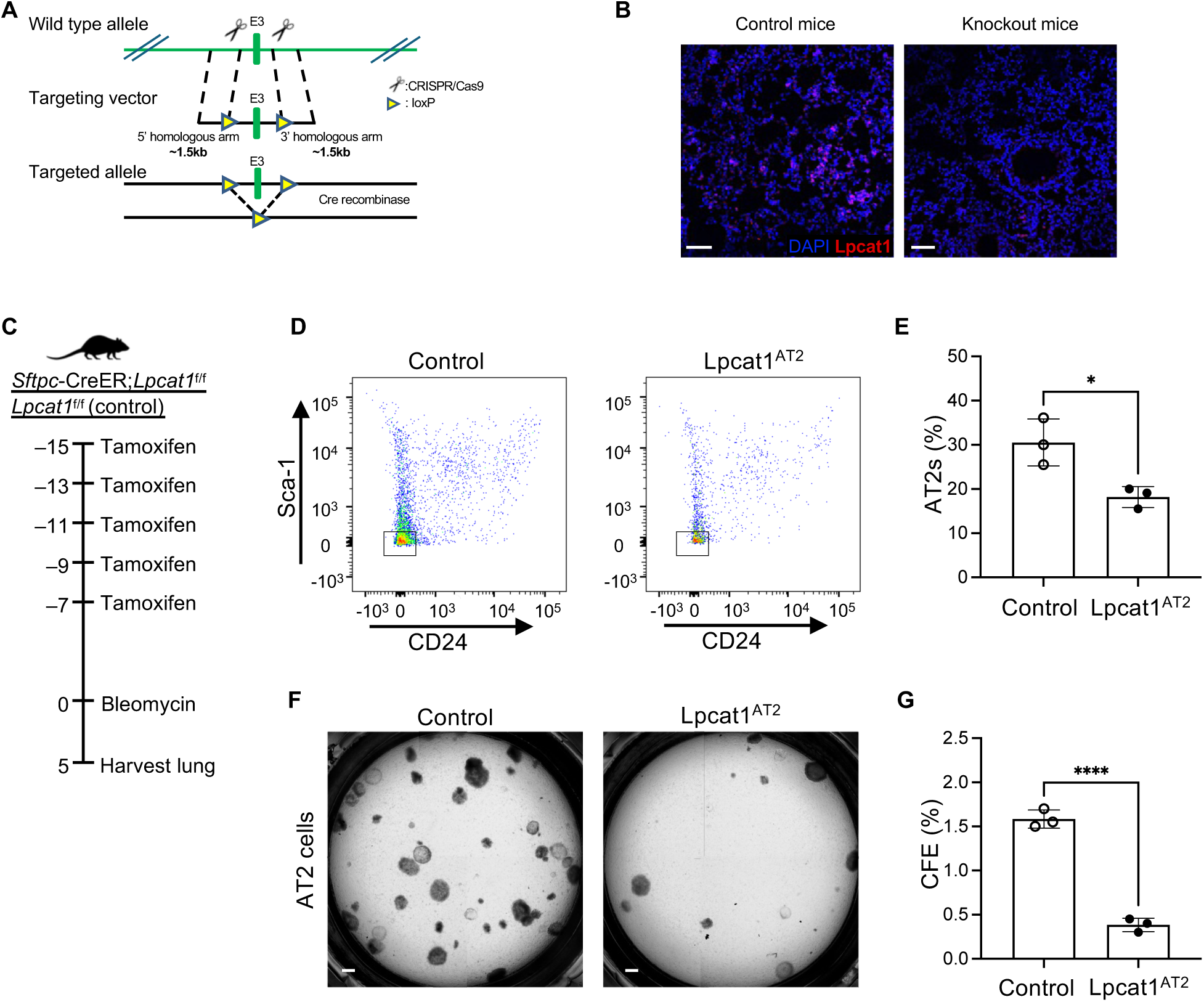
Knockout of Lpcat1 significantly prevents AT2 recovery from bleomycin-injured mouse lungs. (A) Targeting strategy to generate *Lpcat1* floxed mice, utilizing the Cre-loxP system to conditionally knock out exon 3. (B) Immunofluorescence staining of Lpcat1 (red) in lung sections from *Lpcat1* knockout (Lpcat1^AT2^) and control mice. Scale bars: 100 μm. Staining was performed on lung sections from 4 Lpcat1^AT2^ and 4 control mice. (C) Experimental layout of 8-10-week-old Lpcat1^AT2^ and control mice treated with 1.25 U/kg bleomycin following tamoxifen injection, sacrificed at day 5. (D) Flow-cytometry gating strategies for AT2s sorting from Lpcat1^AT2^ and control mice after bleomycin injury. (n = 3 per group, *p < 0.05). (E) Percentage of AT2s (EpCAM^+^CD31^−^CD34^−^CD45^−^CD24^−^Sca-1^−^) in total gated lung epithelial cells per lung. (F) Representative images of 3D organoid cultures (day 14) of AT2s from Lpcat1^AT2^ and control mice after bleomycin injury. Scale bars, 100 μm. (G) CFE of AT2s after 2 weeks of 3D organoid cultures. (n = 3 per group, ****p < 0.0001).

To determine the role of LPCAT1 in regulating AT2 progenitor cell function in vivo, we applied a bleomycin-injured mouse model in *Lpcat1*^AT2^ mice. The mice were administered 1.25 U/Kg bleomycin one week after tamoxifen injection and the lungs were harvested five days post-injury for analysis (Figure 4C). Lung tissues were dissociated into single cells, which were subsequently stained with cell surface markers and analyzed by flow cytometry. AT2s (EpCAM^+^CD24^−^Sca-1^−^) were gated from epithelial cell population (Figure 4D). The analysis revealed a decrease in both the frequency and absolute number of AT2s in the lungs of *Lpcat1*^AT2^ mice (Figure 4E).

To evaluate renewal capacity, AT2s were isolated and analyzed using a 3D organoid culture model. The results indicated that organoid formation was significantly diminished following *Lpcat1* deletion (Figure 4F and 4G). These data suggested that LPCAT1 promotes AT2 cell renewal and plays a protective role in AT2s against bleomycin-induced injury.

### Mice with LPCAT1 deletion in AT2 cells developed spontaneous lung fibrosis

Although LPCAT1 has been suggested to contribute to lung injury (*22, 33, 45*), its function in IPF remains unknown. To investigate this, we examined the long-term consequences of AT2-specific *Lpcat1* deletion on lung fibrosis. 8- to 10-week-old *Lpcat1*^AT2^ mice and littermate controls received five doses of 200 mg/kg tamoxifen, and lung fibrosis was assessed when the mice reached 12 months of age (Figure 5A). The 12-month-old *Lpcat1*^AT2^ mice developed spontaneous lung fibrosis, as evidenced by enhanced trichrome staining (Figure 5B and S3A).

**Figure 5:**
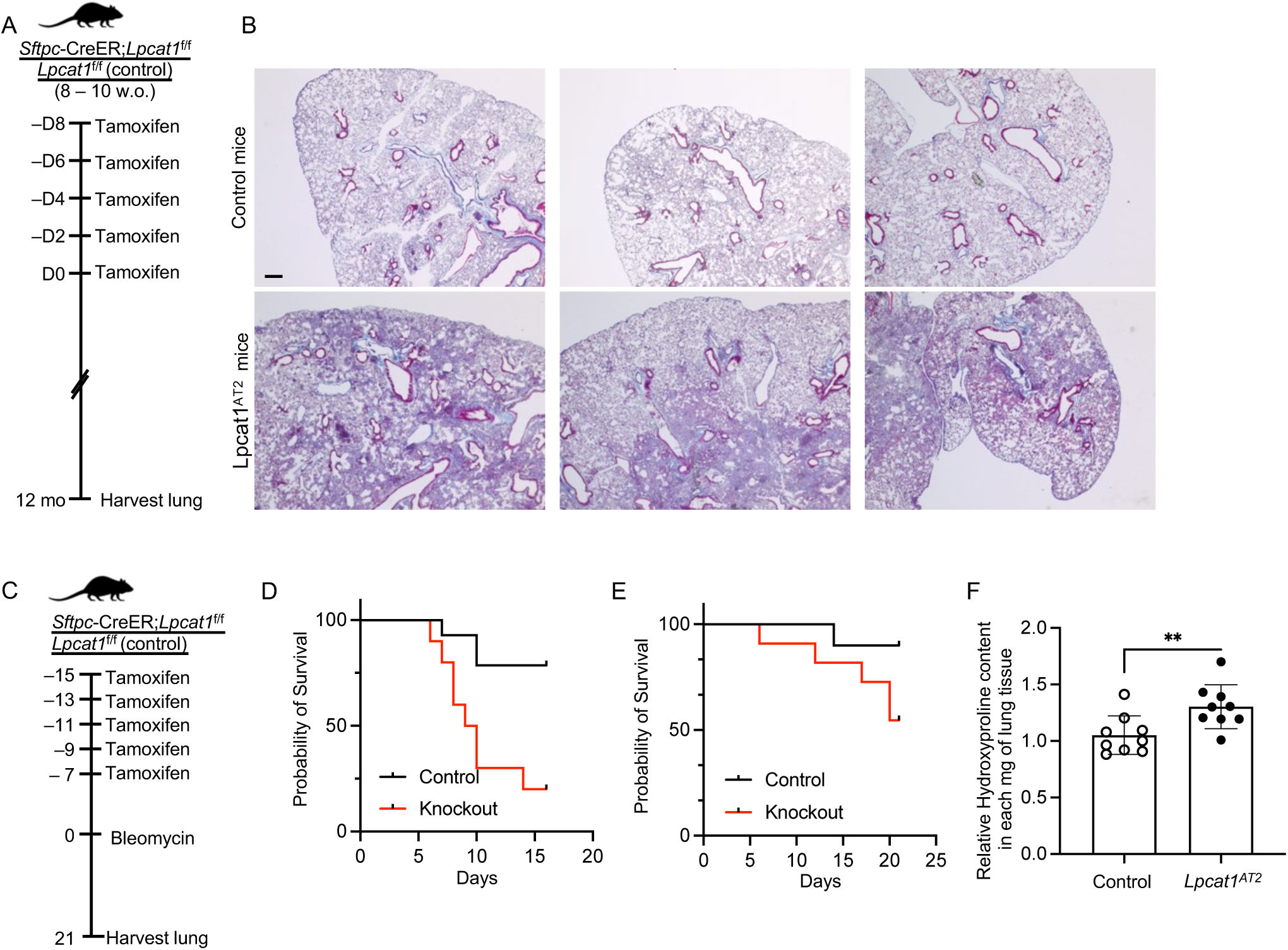
Exaggerated lung fibrosis in Lpcat1^AT2^ mice. (A) Experimental layout: 8 – 10 weeks old mice were given tamoxifen five times. The lungs were harvested when the mice reached to 12 months old. (B) Trichrome staining of lung sections from 12-month-old Lpcat1^AT2^ and control mice. Scale bars: 100 μm. Staining was performed on lung sections from 5 Lpcat1^AT2^ and 5 control mice. (C) Experimental layout: 8 – 10 weeks old mice were treated with 2.5 U/kg or 1.25 U/kg bleomycin following five doses of tamoxifen injections. The lungs were harvested 21 days after bleomycin. (D) Survival percentage of Lpcat1^AT2^ and control mice after 2.5 U/kg bleomycin treatment (** p < 0.01 by Log-rank). (E) Survival percentage of Lpcat1^AT2^ and control mice after 1.25 U/kg bleomycin treatment (ns, not significant by Log-rank). (F) Relative hydroxyproline content in the lungs of Lpcat1^AT2^ (n = 9) and control (n = 9) mice at day 21 after 1.25 U/kg bleomycin treatment (**p < 0.01).

### Mice with *Lpcat1* deletion in AT2 cells were susceptible to bleomycin injury

We next investigated lung fibrosis in *Lpcat1*^AT2^ mice subjected to bleomycin-induced lung injury, to more accurately mimic the characteristics of IPF. The mice were treated with two doses (1.25 or 2.5 U/kg) of bleomycin one weeks after tamoxifen injections and were harvested 21 days post bleomycin injury (Figure 5C). At the low dose, around 45% of *Lpcat1*^AT2^ mice died, showing shortened survival compared with control mice (Figure 5D), while at high dose, most of Lpcat1^AT2^ mice died (Figure 5E).

*Lpcat1*^AT2^ mice also showed increased hydroxyproline content compared with control mice (Figure 5F). These findings together show that deletion of *Lpcat1* in AT2s leads to spontaneous lung fibrosis, increases susceptibility to bleomycin-induced injury, and results in more severe fibrosis.

### Dysregulated phospholipids in LPCAT1-knockout AT2 cells

Since LPCAT1 is a key enzyme regulating PC remodeling in AT2s, and is markedly downregulated in IPF, we hypothesize that the PC content would be altered in IPF AT2s. PC levels were assessed in AT2s deficient for LPCAT1, using the *Lpcat1*^AT2^ mouse model. The results confirmed that total PC content was reduced in AT2s from *Lpcat1*^AT2^ mice (Figure 6A). In AT2s, PC, particularly DPPC, constitutes about 90% of the lipids in pulmonary surfactant. We next collected bronchoalveolar lavage (BAL) from *Lpcat1*^AT2^ mice to assess total PC levels. Our findings indicated that total PC content was significantly decreased following Lpcat1 knockout (Figure 6B).

**Figure 6:**
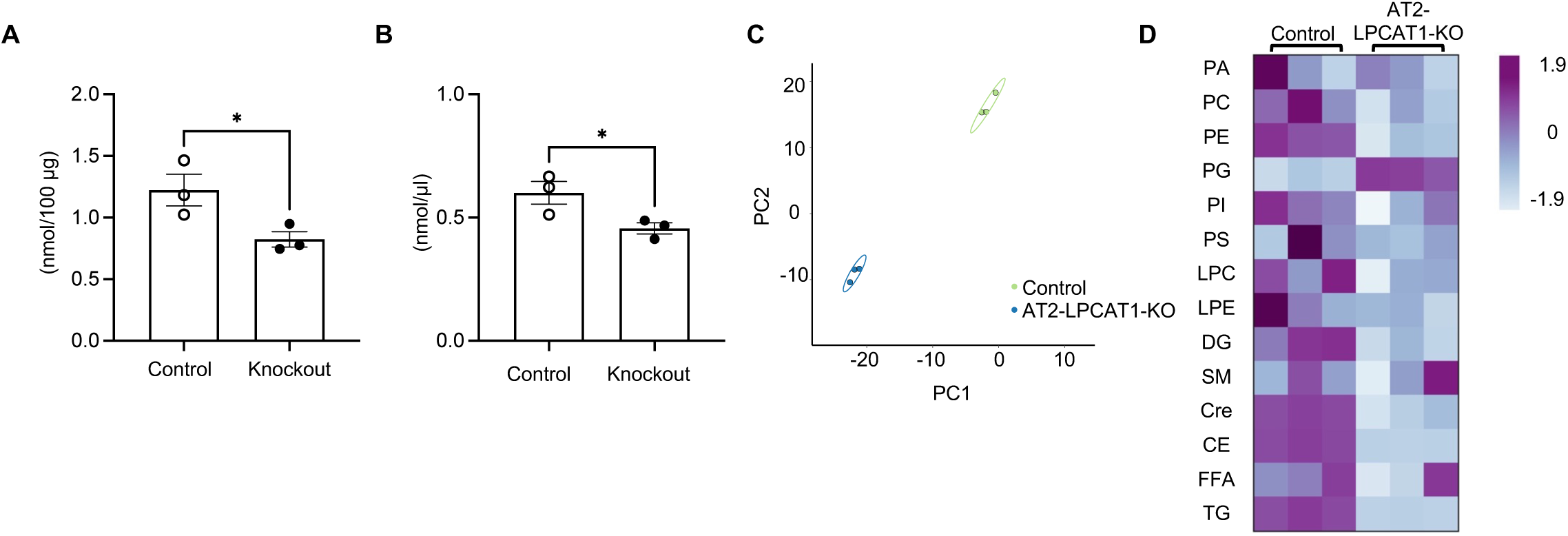
LPCAT1 regulates phospholipid metabolism in AT2 cells. (A) Phosphatidylcholine (PC) content in lung tissue of Lpcat1A^T2^ and control mice (n = 3 per group, *p < 0.05). (B) PC content in BAL of Lpcat1^AT2^ and control mice (n = 3 per group, *p < 0.05). (C) Principal Component Analysis (PCA) plot of lipid components by shotgun lipidomic assay in *LPCAT1*-KO cell lines and healthy AT2 cell (control) (n = 3 per group). (D) Heatmap from the shotgun lipidomic assay highlighting changes in lipid profiles of *LPCAT1*-KO cell lines and healthy AT2 cell (control) (PA = Phosphatidic acid, PC = Phosphocholine, PE = Phosphoethanolamine, PG = Phosphoglycerol, PI = Phosphoinositol, PS = Phosphoserine, LPC = Lysophosphatidylcholine, CE = Cholesterol, Cer = Ceramide, DG = Diglyceride, TG = Triglycerides, SM = Sphingomyelin, FAA = Free fatty acids) (n = 3 per group).

In mammalian cells, PC interacts with other classes of phospholipids. Concurrently, there are interactions among phospholipids, diglycerides, triglycerides, sphingomyelin, and cholesterol, which are vital for numerous physiological and pathophysiological processes (*46, 47*). Although several reports have described changes in lipid metabolism in IPF (*48–50*), no studies have specifically investigated how LPCAT1 influences the interaction of PC with other lipids in IPF AT2s. Using wild-type and LPCAT1 knockout (AT2-LPCAT1-KO) healthy AT2 cell lines, we characterized the lipid profiles of AT2s after LPCAT1 deletion. The shotgun lipidomic assay revealed shifts in lipid components following the LPCAT1 knockout, as evidenced by the principal component analyses (PCA) plots (Figure 6C). The phospholipid profile studies indicated that the levels of PC, phosphatidylethanolamine (PE), phosphatidylinositol (PI), and phosphatidylglycerol (PG) significantly changed after LPCAT1 deletion, suggesting alterations in both cellular and surfactant lipid structures (Figure 6D).

Interestingly, significant changes were also observed in Diglyceride (DG), Ceramide (Cer), Triglyceride (TG), and Cholesterol (CE) (Figure 6D). These findings provide evidence that changes in LPCAT1 expression levels result in a comprehensive shift in the lipid profile of AT2 cells. This altered lipid composition likely influences the functionality of various cellular organelles, including mitochondria, endoplasmic reticulum, and cell membranes.

For further investigation, we conducted Ingenuity Pathway Analysis (IPA) on healthy (AT2-LPCAT1-KO) and IPF (AT2-LPCAT1-OE) cell lines, comparing each to their controls. We observed that several lipid biosynthesis pathways, including PPAR signaling, PPARα/RXRα activation, glycerophospholipid biosynthesis, triglyceride biosynthesis, and diacylglycerol biosynthesis I, were downregulated following LPCAT1 knockout in healthy AT2s (Figure S4A). Conversely, LPCAT1 overexpression in IPF AT2s upregulated these pathways (Figure S4B). Collectively, these findings suggest that LPCAT1 mediates the PC remodeling and is essential for regulating lipid metabolism in AT2s.

### Decreased LPCAT1 led to mitochondria dysfunction in IPF AT2 cells

The structure and function of mitochondria are regulated by the lipid composition of their membranes (*51–53*). PC along with phosphatidylethanolamine (PE) and cardiolipin (CL) are the major components of mitochondrial membrane (*52*). Especially, they contribute to the inner mitochondrial membrane structure and curvature (*52, 53*). Alterations in phospholipids can change the fluidity and rigidity of the inner mitochondrial membrane, thereby compromising mitochondrial function. The lipidomic assay demonstrated that the major lipids of mitochondrial membrane lipids (diglycerides, phospholipids, and ceramide) were altered following LPCAT1 deletion. We suggest that these changes contribute to the loss of mitochondrial function in AT2s, leading to AT2 dysfunction in IPF. Additionally, IPA analysis indicated downregulation of the mitochondrial dysfunction pathway in healthy AT2s after LPCAT1 deletion (Figure S4A). In contrast, LPCAT1 overexpression in IPF AT2s led to upregulation of these pathways (Figure S4B). We suggested that these changes contribute to the loss of mitochondrial function in AT2s, leading to AT2 dysfunction in IPF.

Mitochondrial function in IPF and healthy AT2 cell lines was assessed by measuring the oxygen consumption rate (OCR) in real time with a Seahorse metabolic analyzer. The result indicated that OCR was significantly lower in IPF AT2s compared to healthy AT2s (Figure 7A). These changes were observed across all phases of mitochondrial respiration, including basal respiration, ATP production, proton leak, maximal respiration, and spare respiratory capacity (Figure 7B).

**Figure 7:**
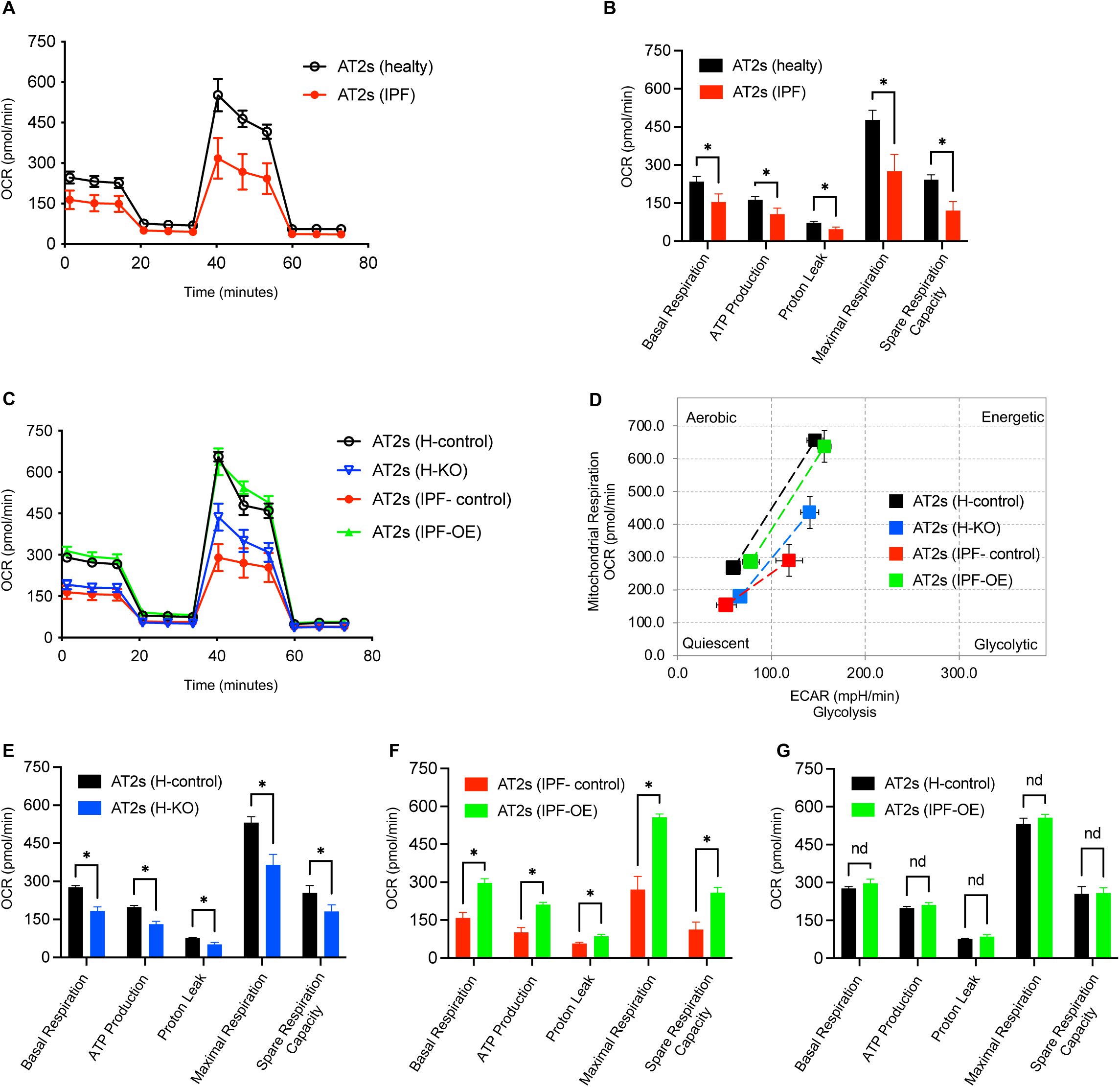
Effect of LPCAT1 on mitochondrial function. (A) Changes in oxygen consumption rate (OCR) in healthy and IPF AT2 cells using the Seahorse Respirometry XFe96. (B) Mitochondrial respiration changes in healthy and IPF AT2 cells, analyzed for basal respiration, ATP production, proton leak, maximal respiration, and spare respiratory capacity. (C) OCR in AT2-*LPCAT1*-KO and AT2-*LPCAT1*-OE cell lines and their respective control cell lines. (D) XF Cell Energy Phenotype in AT2-*LPCAT1*-KO and AT2-*LPCAT1*-OE cell lines and their respective control cell lines. (E-G) Mitochondrial respiration changes in AT2-*LPCAT1*-KO and AT2-*LPCAT1*-OE cell lines and their respective control cell lines, analyzed for basal respiration, ATP production, proton leak, maximal respiration, and spare respiratory capacity. Data are presented as the mean ± SEM from 6 replicate wells. *p < 0.05 by unpaired t-tests.

Furthermore, to explore the function of LPCAT1 in regulating OCR, Seahorse assay was conducted on healthy (AT2-LPCAT1-KO) and IPF (AT2-LPCAT1-OE) cell lines. Deletion of LPCAT1 in healthy AT2 cell lines resulted in a significant decrease in OCR compared to control cells. In contrast, LPCAT1 overexpression in IPF AT2 cells led to a significant elevation of OCR compared to control cells (Figure 7C). The XF cell energy phenotype test showed that LPCAT1 overexpression in IPF AT2 cells produced an energy phenotype closely resembling that of healthy AT2 cells. On the other hand, LPCAT1 deletion in healthy AT2s shifted their energy phenotype to resemble that of IPF AT2s (Figure 7D).

Comparison of healthy (AT2-LPCAT1-KO) and IPF (AT2-LPCAT1-OE) cell lines with their respective controls revealed significant changes in mitochondrial respiration throughout all phases (Figure 7E and 7F). Notably, no differences in mitochondrial respiration phases were observed when comparing IPF (AT2-LPCAT1-OE) cell lines to healthy AT2 cells (Figure 7G), suggesting that mitochondrial function might be restored by activating LPCAT1 in IPF AT2 cells.

### Expression-based drug screen identified compounds that restore LPCAT1-mediated function

Together, these findings suggest that reactivation of LPCAT1 in IPF AT2s may enhance cell function and potentially reduce lung fibrosis. To this end, we established an expression-based high-content screening system (*54–56*) to identify candidate compounds capable of upregulating LPCAT1 expression. We engineered an LPCAT1 knock-in cell line tagged with a Luciferase-mCherry fusion allele and performed drug screening (Figure S5A). Specifically, an IRES-Luciferase-mCherry fusion cassette was inserted immediately after the STOP codon of the *LPCAT1* gene using CRISPR-Cas9 (Figure S5B). To facilitate detection of increased expression during screening, we selected A549 cells, as they exhibit relatively low basal LPCAT1 expression compared to other lung cell lines (Figure S5C). After introducing the LPCAT1 knock-in construct into A549 cells, single mCherry-positive clones with low expression were isolated and expanded for subsequent drug screening (Figure S5D). The screening was performed using two libraries: LOPAC1280, a well-established collection of Pharmacologically Active Compounds, and NPWII, comprising structurally diverse FDA-approved drugs across a broad range of therapeutic areas (*57–60*). Both libraries were prepared and maintained by Molecular Screening Shared Resource at UCLA.

After the initial screening, 35 hits were further subjected to dose-response assays. We chose two compounds – the antimalarial Artesunate and the PLA2 inhibitor ONO-RS-082 – for further analysis (Figure 8A). Antimalarial drug Artesunate has been shown to exert anti-fibrotic effects in cardiac fibrosis (*61, 62*). ONO-RS-082 is a reversible PLA2 inhibitor (*63, 64*) that has been tested in patients with pulmonary arterial hypertension (*65*). To confirm the upregulation of LPCAT1 gene expression, we cultured cells in the presence of ONO-RS-082 and Artesunate. Both compounds significantly increased *LPCAT1* levels in AT2 cell lines (Figure 8B). Building on our previous findings that LPCAT1 regulates AT2 renewal and recovery, we treated AT2 cells from both healthy and injured mice with these compounds. We found that Artesunate enhanced CFE only in healthy AT2 cells (Figure 8C and 8D), while ONO-RS-082 enhanced CFE exclusively in injured AT2 cells (Figure 8E and 8F). The interaction between PLA2 and LPCAT1 in AT2 cells is well-established within the PC remodeling cycle (*25*). Our results suggest that the PLA2 inhibitor promotes AT2 recovery by directly upregulating deficient LPCAT1, but this effect is absent when LPCAT1 levels are normal. The interactions between Artesunate and LPCAT1 in AT2 cells remain unclear, although the results suggest that Artesunate may elevate LPCAT1 levels through an indirect mechanism. Collectively, these findings indicate that these compounds may improve IPF outcomes. Therefore, we treated a bleomycin-induced mouse model with Artesunate or ONO-RS-082 as shown in Figure 8G. Interestingly, Artesunate significantly reduced hydroxyproline content after bleomycin treatment compared to control mice (Figure 8H) and improved survival rates (Figure 8I). The mechanisms by which these drugs upregulate LPCAT1 expression are still under investigation. Nevertheless, our findings open new avenues for developing potential therapies for lung fibrosis.

**Figure 8.**
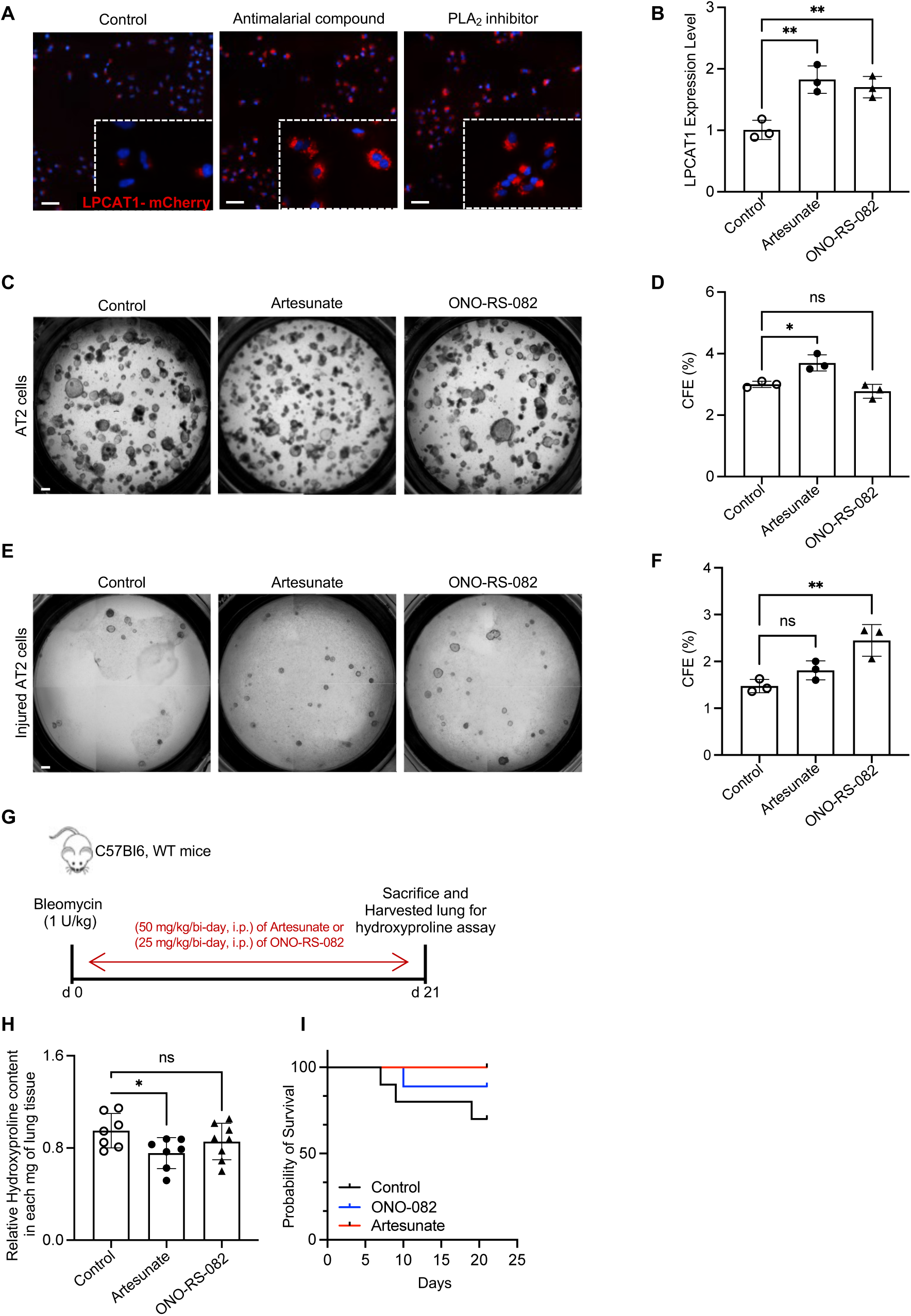
Drug screening identified compounds that upregulate LPCAT1 expression. (A) Drug screening identified antimalarial compound Artesunate and PLA2 inhibitor ONO-RS-082. Scale bars, 100 μm. (B) qPCR analysis of *LPCAT1* expression in healthy AT2 cell lines cultured with Artesunate and ONO-RS-082 (n = 3 per group, ns, no significant, *p < 0.05). (C and D) Representative images of 3D organoid cultures (day 14) and CFE of mouse AT2 cells in the presence of Artesunate or ONO-RS-082 (n = 3 per group, ns, no significant, *p < 0.05). (E and F) Representative images of 3D organoid cultures (day 12) and CFE of injured mouse AT2 cells in the presence of Artesunate or ONO-RS-082 (n = 3 per group, ns, no significant, **p < 0.01). (G) Experiment layout. (H) Relative hydroxyproline content in the lungs of mice treated with Artesunate (n = 7), ONO-RS-082 (n = 8), and control (n = 7) (ns, no significant, *p < 0.05). (I) Survival curves of mice treated with Artesunate or ONO-RS-082 after 1 U/kg bleomycin treatment (ns, not significant by Log-rank).

## Discussion

Our previous work has shown that impaired AT2 progenitor renewal plays a critical role in the development of lung fibrosis (5, 11, 26). However, the molecular mechanisms driving AT2 cell dysfunction in IPF remain poorly understood. Metabolic studies have linked dysregulated lipid levels to the pathogenesis of IPF (*66, 67*). In this study, we identified LPCAT1 as a critical regulator of AT2 progenitor renewal and lipid metabolism in IPF. We hypothesized that LPCAT1 deficiency disrupts PC homeostasis and mitochondrial function, thereby impairing AT2 renewal and promoting lung fibrosis. To test this hypothesis, we employed genetic approaches to generate cell lines with LPCAT1 knockout or overexpression and confirmed that LPCAT1 regulates PC metabolism, mitochondrial function, and AT2 renewal in 3D organoid assays. We further generated a mouse line with AT2 cell–specific deletion of *Lpcat1* and demonstrated that LPCAT1 regulates PC metabolism, AT2 renewal in organoid assays, and, importantly, lung fibrosis in vivo. Through expression-based drug screening, we identified phospholipase A2 (PLA2) inhibitors and antimalarials as compounds that upregulate LPCAT1, with artesunate significantly reducing fibrosis in vivo. Collectively, these findings establish LPCAT1 as a central mediator of AT2 progenitor function and a promising therapeutic target for IPF.

The PC serves as the main phospholipids in AT2s (*15, 18*). It is considered essential for maintaining cell architecture and regulating cell function and the production of pulmonary surfactants (*22, 24, 27*). We applied the lipidomic shotgun approach to analyze the lipid profile in AT2 cell lines after knocking out *LPCAT1*. The findings indicated that phospholipids significantly changed after knocking out *LPCAT1* in AT2s, particularly in the classes of PC, PE, PI, and PG, which are considered the primary lipid components of pulmonary surfactants and are involved in cellular function. Additionally, changes were noted in DG, Cer, TG, and CE, which are associated with the function of cellular organelles, such as mitochondria. In addition, the pathway analysis of healthy (AT2-LPCAT1-KO) AT2 cell lines compared to their control revealed a downregulation in several lipid biosynthesis pathways including PPAR signaling, PPARa/RXRa activation, glycerophospholipid biosynthesis, triglyceride biosynthesis, and diacylglycerol biosynthesis I. Conversely, the overexpression of LPCAT1 in the IPF AT2 cell lines resulted in the upregulation of these pathways. Our lipidomic and pathway studies suggest that the PC remodeling mediated by LPCAT1 interacts with other lipids to maintain the lipid profile of AT2 cells. Notably, a compelling study demonstrated that the targeted inhibition of LPCAT1 degradation during inflammatory lung injury represents a viable therapeutic strategy for elevating LPCAT1 levels, consequently preserving surfactant biosynthesis and promoting pulmonary function (*68*). Evidence underscores the critical role of LPCAT1 in maintaining balanced PC biosynthesis in lung epithelial cells, through the interplay of the CDP-choline and de novo pathways (*69, 70*). Furthermore, an in vivo study revealed that LPCAT1 plays an essential role in surfactant production in mouse AT2 cells, critically contributing to PC synthesis necessary for the successful transition to air breathing (*23*). A lipidomic analysis of BAL from mice with bleomycin-induced lung fibrosis revealed significant alterations in the lipid profile, primarily characterized by changes in PE, PC, and PG (*71*). These studies are consistent with our findings on the essential role of LPCAT1 in maintaining phospholipid homeostasis, especially PC/LPC remodeling in AT2 cells.

The function and structure of mitochondria are maintained by lipid composition (*51, 72*). Phospholipids in mitochondrial membranes may play a role in the pathogenesis of IPF (*73*). The function of mitochondria is regulated by LPCAT1 (*74*). We performed Seahorse assays to demonstrate the role of LPCAT1 in regulating mitochondrial function in AT2 cells. Our data indicated that IPF AT2 cell lines exhibited mitochondrial dysfunction compared to healthy AT2 cell lines. Furthermore, *LPCAT1* overexpression in IPF AT2 cell lines restored mitochondrial function and energy phenotype of the cells. PC, PE, and cardiolipin (CL) are the primary constituents of the mitochondrial membrane (*52, 75*). Since LPCAT1 is pivotal in surfactant phospholipid synthesis (*23, 34*), and in phospholipid metabolism (*69, 70*), primarily through the regulation of PC levels in cells. Given its role in phospholipid regulation, it is likely that LPCAT1 also influences CL levels in mitochondria. Studies indicate that abnormalities in cardiolipin (CL) may play a pathological role in lung inflammation and fibrosis (*76, 77*).

Studies on LPCAT1-deficient mice have shown that the physical properties of pulmonary surfactants are altered due to changes in PC content in the lungs (*22*). The critical role of LPCAT1 in the biogenesis of membrane phospholipids and surfactant lipids in alveolar AT2 cells has been reported (*33, 35*). *Lpcat1* gene-trap mice exhibited a significant reduction in surfactant protein levels and markedly decreased activity in newborns (*23*). To explore the role of LPCAT1 in the development of IPF, we generated *Lpcat1*^AT2^ mouse model. Our findings clearly indicated that the renewal capacity of freshly isolated AT2 cells from these mice was significantly reduced following the deletion of *Lpcat1*. Interestingly, the absence of *Lpcat1* in AT2s resulted in spontaneous lung fibrosis as the mice aged. We observed an increase in lung fibrosis and mortality rates after lung injury, accompanied by a loss in the recovery ability of AT2s. Dysregulated lipid metabolism and impaired surfactant homeostasis have been observed in lung fibrosis (*24, 78*). Loss of progenitor function of AT2 cells and senescence play key roles in the pathogenesis of lung fibrosis (*5, 79*). To our knowledge, these are the first data demonstrating that LPCAT1 is essential for maintaining the proper progenitor function of AT2s and is involved in the progression of pulmonary fibrosis.

Unfortunately, no specific drugs or activators have been identified that directly upregulate LPCAT1 expression. We then designed an expression-based high-content screen system (*54–56*) to screen the Library of Pharmacologically Active Compounds (LOPAC1280) and the FDA-approved NPWII library (*57–60*) to identify compounds that potentially increase LPCAT1 expression. We identified PLA2 inhibitors and antimalarial drugs. Antimalarial artesunate has been shown to be anti-fibrotic in cardiac fibrosis (*61, 62*) and in bleomycin-induced lung fibrosis in rodents (*80, 81*). ONO-RS-082 is a reversible PLA2 inhibitor (*63, 64*) and has been tested in patients with pulmonary arterial hypertension (*65*). PLA2 has been shown to regulate lung injury and inflammation (*82*). We confirmed both antimalarial artesunate and PLA2 inhibitor ONO-RS-082 increased LPCAT1 expression. We further demonstrated that PLA2 inhibitor ONO-RS-082 was unable to change outcome of lung fibrosis, but antimalarial artesunate attenuated bleomycin-induced lung fibrosis in mice in vivo. The mechanisms these compounds regulate LPCAT1 expression are under active investigation.

In summary, this study demonstrates that LPCAT1 expression is significantly reduced in AT2 cells during lung fibrosis. We elucidated the role of LPCAT1 in regulating AT2 cell renewal and the progression of pulmonary fibrosis, both in vivo and in vitro. Notably, alterations in LPCAT1 expression disrupted the lipid profile of AT2 cells. Furthermore, we identified mitochondrial dysfunction in IPF AT2s as a consequence of LPCAT1 downregulation. Our findings indicate that promoting LPCAT1 expression in IPF AT2s restores phosphatidylcholine metabolism and mitochondrial function, offering a potential strategy to enhance AT2 recovery and alleviate fibrosis in patients with IPF.

## Materials and Methods

### Donor lung tissues

The use of human tissues for research was approved by the Institutional Review Boards of Cedars-Sinai Medical Center (CR00005442), with informed consent obtained from all participants. The study involved healthy donors and IPF patients, including both males and females.

### Mice

All animal experiments were approved by the Institutional Animal Care and Use Committee at Cedars-Sinai Medical Center (Protocol IACUC008529). All mice were housed in a pathogen-free facility at Cedars-Sinai Medical Center. *Lpcat1*^floxed^ mice were created by introducing loxP sites flanking exon 3 via CRISPR/Cas9 system. *Lpcat1*^floxed^ mice were crossed with *Sftpc*-CreER mice (*12*), generating *Sftpc*-CreER;*Lpcat1*^flox/flox^ mice (*Lpcat1*^AT2^). Wild-type C57Bl/6J mice were obtained from The Jackson Laboratory. All mice were backcrossed onto the C57Bl/6J background for more than six generations. Mice were used at 8–12 weeks. They were randomly assigned to treatment groups, matched by age and sex.

### Human lung dissociation and AT2 cells isolation

Human lung tissue and AT2 cells isolation was applied as previously described (*6*). Briefly, lung tissues were minced and digested with 2 mg/ml Dispase II (Sigma), followed by 10 U/ml elastase (Worthington Biochemical Corp.) and 100 U/ml DNase I (Sigma). Cells were then filtered through a 100 μm cell strainer and subjected to red blood cell lysis (BioLegend) to obtain a single-cell suspension. Antibodies used included anti-human CD31 (clone WM59, Catalog # 303118), anti-human CD45 (clone HI30, Catalog # 304016), and anti-human EpCAM (clone 9C4, Catalog # 324212) from BioLegend, and anti-HTII-280 (Terrace Biotech, Catalog # TB-27AHT2-280). AT2 cells were sorted as EpCAM^+^HTII-280^+^CD31^−^CD45^−^ using a BD Symphony S6 Cell Sorter.

### Mouse lung dissociation and AT2 cells isolation

Mouse lung tissue dissociation and AT2 cell isolation were previously described (*6*). Briefly, lungs were perfused with 10 mL PBS, then digested with 4 U/mL elastase (Worthington Biochemical Corp.) and 100 U/mL DNase I (Sigma). Cells were filtered through a 100 μm strainer, subjected to red blood cell lysis (BioLegend), and resuspended as a single-cell suspension. The suspension was incubated with antibodies: PE/Cyanine7 anti-mouse CD326 (Ep-CAM) (BioLegend, Clone G8.8, #118215), FITC anti-mouse CD24 (BioLegend, Clone M1/69, #101806), APC anti-mouse Ly-6A/E (Sca-1) (eBioscience, Clone D7, #17598181), PE anti-mouse CD31 (BioLegend, Clone MEC13.3, #102507), PE anti-mouse CD34 (BioLegend, Clone HM34, #128609), and PE anti-mouse CD45 (BioLegend, Clone 30-F11, #103105). Dead cells were excluded using 7-AAD (BD Biosciences). AT2 cells were sorted as EpCam^+^CD31^-^CD34^-^CD45^-^CD24^-^Sca-1^-^ using a BD Symphony S6 Cell Sorter.

### 3D organoid cultures of AT2 cells

As previously described (*5*), 3D organoid cultures were established using flow-sorted human (EpCAM^+^HTII-280^+^) or mouse (EpCAM^+^CD24^−^Sca-1^−^) AT2 cells. These were cultured with MLg2908 lung fibroblasts (ATCC, CCL-206) in a 1:1 mix of Matrigel (Growth Factor Reduced Basement Membrane Matrix, catalog 354230) and medium. A total of 3 × 10³ AT2 cells and 2 × 10⁵ MLg2908 cells were seeded in 100 μl of the Matrigel/medium mix onto 24-well, 0.4 μm Transwell inserts, with 410 μl of medium in the lower chamber. The medium consisted of DMEM/F12 (Gibco, Catalog 11320-033) supplemented with 10% FBS (Invitrogen), 1x insulin/transferrin/selenium (Gibco), 100 U/ml penicillin, 100 μg/ml streptomycin, and 10 µM SB431542 (Sigma). Cultures were maintained in medium alone or with specified inhibitors or treatments: 1, 3, 10, or 20 µM TSI-01 (Cayman Chemical, Catalog 17628), 5 µM Artesunate (Cayman Chemical, Catalog 11817; Sigma, 704878-75), or 5 µM ONO-RS-082 (Cayman Chemical, Catalog 20243; Sigma, O0766). AT2 cell lines were cultured in a feeder-free 3D organoid culture as previously detailed (*83*). Briefly, immortalized AT2 cells were resuspended in Matrigel and plated into a Matrigel-coated 24-well plate in domes (5 × 10^3^ cells in 50 μl Matrigel/well). Cells were cultured under serum-free conditions consisting of DMEM/F12, supplemented with 1 × B-27 without vitamin A (Gibco, Catalog 12587010), 100 U/ml penicillin, 100 μg/ml streptomycin, 20 ng/ml human recombinant EGF (ThermoFisher, Catalog PHG0311), 20 ng/ml human recombinant FGF10 (#100-26, Peprotech), and 10 μM Y-27632 (Enzo Life Sciences, Catalog 270–333). The medium was changed every 3 days alone or with specified inhibitors or treatments according to experiment. Colony formation efficiency (CFE%) was calculated as (number of organoids formed)/(number of cells seeded) × 100. The cell growth rate was measured by IncuCyte S3 Live Cell Analysis System (Sartorius).

### Generation of immortalized AT2 cell lines

Immortalized AT2 cell lines were generated using a modified protocol from Dr. Borok and Offringa’s group (*44*). Briefly, EpCAM^+^ cells were magnetically enriched from healthy and IPF lung tissues using CD326 (EpCAM) microbeads (Miltenyi Biotec, catalog 130-061-101). Cells were resuspended in a 3:1 mix of DMEM/F12 (Corning, Catalog 10-090-CV) and DMEM (Gibco, Catalog 21063-029), supplemented with 5% FBS, 0.4 mg/mL hydrocortisone (Sigma-Aldrich, Catalog H0888), 5 mg/mL insulin (Sigma-Aldrich, Catalog I0516), 8.4 ng/mL cholera toxin (Sigma-Aldrich, Catalog C8052), 10 ng/mL human recombinant EGF, Antibiotic-Antimycotic (Gibco, Catalog 15240-062), and 10 μM Y-27632. Cells were allowed to adhere for two days, with media changed every 2–3 days. At passage 3–4, cells were infected with SV40 Large T antigen lentivirus (customized from VectorBuilder) for immortalization, followed by selection with G418 (Sigma-Aldrich, Catalog G8168). AT2 cells were sorted as EpCAM^+^HTII-280^+^CD31^-^CD45^-^ from total epithelial cells.

### Generation of LPCAT1-KO *and* LPCAT1-OE cell lines

To generate LPCAT1 knock-out (LPCAT1-KO) and LPCAT1 overexpression (LPCAT1-OE) cell lines using CRISPR-Cas9 system, immortalized AT2 cells from healthy lungs were infected with a commercial LPCAT1 sgRNA CRISPR/Cas9 All-in-One Lentivirus (Abm, Catalog #27555111) for knockout. For overexpression, immortalized AT2 cells from IPF lungs were infected with a commercial LPCAT1 CRISPR sgRNA lentivirus (Abm, Catalog #27555121).

### Generation of LPCAT1-KI cell line

To generate a stable LPCAT1 knock-in cell line, A549 cell line (ATCC, CCL-185) was transduced with a gRNA/Cas9 system targeting the locus immediately downstream of the LPCAT1 stop codon, along with a donor vector carrying a luciferase-mCherry cassette (customized from VectorBuilder). Specifically, the luciferase-mCherry cassette was inserted downstream of the LPCAT1 stop codon via a bicistronic IRES element. In this configuration, mCherry fluorescence or luciferase luminescence serves as a real-time reporter of endogenous LPCAT1 promoter activity, thereby reflecting endogenous LPCAT1 expression.

### scRNA-seq and data collection

The scRNA-seq datasets GSE157997 and GSE157995, generated by our group (*5*) at the Cedars-Sinai Medical Center Genomic Core, along with datasets GSE132915 (35), GSE150068 (36), GSE132771 (37), and GSE128033 (*19*) from other groups, were analyzed to assess LPCATs expression in healthy and IPF human samples, as well as in healthy and bleomycin-injured mouse models.

### Bulk RNA-seq and Ingenuity Pathway Analysis

RNA was extracted from AT2-LPCAT1-KO and AT2-LPCAT1-OE cell lines, cultured in 60 mm plates, using the RNeasy Mini Kit (Qiagen). The total RNA was submitted to the UCLA Technology Center for Genomics and Bioinformatics (TCGB) for library preparation and RNA sequencing, performed on a NovaSeq X Plus platform. Differentially expressed genes compared to controls were identified, and the data were analyzed using IPA software (QIAGEN) to investigate canonical pathways.

### Bleomycin instillation

As previously described (*6*), bleomycin instillation was performed on anesthetized mice by surgically exposing the trachea. A 25-G needle was inserted between the tracheal cartilaginous rings to instill 1, 1.25, or 2.5 U/kg bleomycin (Hospira) in 25 μl PBS. Control mice received saline alone. The tracheostomy site was sutured, and mice were allowed to recover. At various time points, mice were euthanized, and lung tissue was collected for experiments.

### Drug treatment

The compounds Artesunate and ONO-RS-082 were dissolved in DMSO and diluted with PBS before administration in mice. Starting on day 1 post-bleomycin treatment, mice received intraperitoneal injections every other day of either 50 mg/kg Artesunate, or 25 mg/kg ONO-RS-082 for 21 days.

### Hydroxyproline assay

Collagen content in mouse lungs was quantified using a standard hydroxyproline method (*84*). Briefly, lung tissues were vacuum-dried and hydrolyzed with 6N hydrochloric acid at 120°C overnight. Hydroxyproline levels were measured and reported as mg per lung. The assay’s ability to fully hydrolyze and accurately recover hydroxyproline from collagen was validated using samples with known quantities of purified collagen.

### Phosphatidylcholine (PC) assay

Total phosphatidylcholine (PC) content in AT2 cell lines, mouse AT2 cells, or bronchoalveolar lavage (BAL) was measured using the Phosphatidylcholine Assay Kit (Abcam, Shanghai, Catalog #ab83377) according to the manufacturer’s protocol. Briefly, BAL collection was performed as previously described (*85*). Mice were anesthetized, and the lungs and heart were surgically exposed. The trachea was cannulated, and the lungs were lavaged three times with 0.8-mL aliquots of cold PBS and assayed directly. For AT2 cell lines and mouse AT2 cells, cells were harvested, washed with PBS, resuspended in assay buffer, and homogenized. After centrifugation, the supernatant was combined with assay buffer, PC hydrolysis enzyme, PC development mix, and OxiRe probe. PC concentration was measured by absorbance at 570 nm using a microplate reader (BioTek, Synergy Neo2).

### Shotgun lipid analysis

Lipidomic analysis was performed at the Lipidomics Core Facility of UCLA following a standardized protocol (*86*). A total of 2 × 10^6^ cells from both AT2 and AT2-LPCAT1-KO cell lines were carefully collected and transferred into serum-free PBS using specialized collection tubes. For shotgun lipidomic analysis, samples were subjected to a dual lipid extraction process, supported by a comprehensive quality assurance panel of 70 lipid standards from Sciex and Avanti Polar Lipids, covering 17 lipid subclasses. Lipid species were quantified using a Sciex 5500 instrument equipped with a differential mobility device Lipidyzer and calibrated with Equi-SPLASH LIPIDOMIX standards for optimal accuracy. Principal Component Analysis (PCA) plots show the percentage of the total variance of PC1 and PC2 were generated using Clustvis.

### Seahorse assays

The Seahorse XF Cell Mito Stress Test (Agilent Technologies, 103015-100) was used to measure oxygen consumption rate (OCR) in cells according to the manufacturer’s protocol. Briefly, 1 × 10⁴ AT2 cell lines were plated in Seahorse XF microplates. One hour before the assay, the culture medium was replaced with Seahorse XF assay medium (Agilent Technologies, 103575-100), and plates were incubated at 37°C in a CO₂-free environment. The mitochondrial stress test was performed under basal conditions, followed by sequential injections of 1.5 μM oligomycin, 0.5 μM FCCP, and 0.5 μM rotenone/antimycin A.

### Expression-based high-content screening

High Throughput Screening was performed at the Molecular Screening Shared Resource of UCLA (*60*). The Library of Pharmacologically Active Compounds (LOPAC1280), a widely utilized collection of bioactive compounds, alongside the NPWII library, which comprises structurally diverse FDA-approved drugs spanning a broad range of therapeutic areas, were screened. 2,000 LPCAT1-KI cells were plated into 384-well plates. Using Echo 555 each well received 500 nM of each compound in 0.5% DMSO, followed by a 48-hour culture period. High-throughput imaging was performed using the Molecular Devices ImageXpress Confocal microscope. Dose-response testing of hit compounds was conducted at concentrations ranging from 50 nM to 5 μM. The number of cells seeded, DMSO toxicity, and culture period were determined using a primary optimization assay.

### Western blotting

Total protein was extracted from cells using RIPA lysis buffer supplemented with protease and phosphatase inhibitors. Protein lysates were homogenized by vortexing, cleared by centrifugation and protein concentration was measured using the BSA protein assay (Thermo Fisher Scientific). Then, Proteins were assessed with western blotting as previously described (*87*). Membranes were incubated with primary antibodies against LPCAT1 (Novus Biologicals, Catalog # NBP1-88923, 1:1000). β-actin (Sigma, Clone AC-74, Catalog # A5316, 1:5000) or GAPDH (Cell Signaling Technology, Clone 14C10, Catalog #2118s, 1:5000) were used as a loading control. Secondary antibodies included anti-rabbit IgG, HRP-linked (Cell Signaling Technology, Catalog #7074), anti-mouse IgG, HRP-linked (Cell Signaling Technology, Catalog #7076), and Peroxidase anti-Rabbit IgG (H+L) (Jackson ImmunoResearch, Catalog #711-036-152). Western blot bands were developed using ECL Western Blotting Substrate (Thermo Fisher Scientific) and visualized with the ChemiDoc MP Imaging System (Bio-Rad). Western blot densitometry was quantified using ImageJ2 software, with band intensities normalized to β-actin or GAPDH.

### Real-Time Quantitative PCR (qPCR)

RNA was isolated using a RNeasy Mini Kit (Qiagen) according to the manufacturer’s instructions. cDNA was prepared using a SuperScript VILO cDNA Synthesis Kit (Thermo Fisher Scientific). Real-time PCR was performed using Power SYBR Green PCR Master Mix (Applied Biosystems) on the ABI 7500 Fast Real-Time PCR System (Applied Biosystems). The specific primers are listed below: Human LPCAT1 forward ACCTATTCCGAGCCATTGACC, and reverse CCTAATCCAGCTTCTTGCGAA. Mouse *Lpcat1* forward CACGAGCTGCGACTGAGC, and reverse ATGAAAGCAGCGAACAGGAG. Human *GAPDH*forward CCCATGTTCGTCATGGGTGT, and reverse TGGTCATGAGTCCTTCCACGATA. Mouse *Gapdh* forward GTGTTCTACCCCCAATGTG, and reverse AAGTCGCAGGAGACAACCTG. Relative gene expression was calculated using the ΔΔCt method and normalization to two housekeeping genes.

### Histology and immunofluorescence staining

Lung tissues from humans or mice, freshly dissected, were fixed in 10% neutral formalin (Sigma-Aldrich) overnight and processed for paraffin embedding using standard methods. Paraffin sections (5 μm) were cut, deparaffinized in xylene, and rehydrated. For histological analysis, sections were stained with hematoxylin and eosin or trichrome. For immunohistochemistry, antigen retrieval was performed using citrate- or Tris-based unmasking solutions prior to antibody staining. Immunofluorescence involved overnight incubation at 4°C with primary antibodies: anti-HTII-280 (Terrace Biotech, #TB-27AHT2-280, 1:50), anti-LPCAT1 (Novus Biologicals, #NBP1-88923, 1:100; Proteintech, #16112-1-AP, 1:100), or anti-SFTPC (Proteintech, #10774-1-AP, 1:200). After washing, sections were incubated with secondary antibodies for 2 h at room temperature: Cy3 goat anti-rabbit IgG (Jackson Immunoresearch, #111-165-003), AF488 goat anti-mouse IgG/IgM (Invitrogen, #A-10680), AF594 goat anti-rabbit IgG (Invitrogen, #A-11012), or AF488 donkey anti-rabbit IgG (Jackson Immunoresearch, #711-545-152). Sections were then washed, stained with DAPI for 10 min, and mounted. Fluorescence was imaged using a Zeiss LSM 780 Confocal Microscope (ZEISS). Slides were quantified with ImageJ2 software.

### Statistics

The statistical difference between groups in the bioinformatics analysis was calculated using the Wilcoxon Signed-rank test. Statistical analysis was conducted using GraphPad Prism software (version 10.4.2) (GraphPad Software, San Diego, CA). Data are expressed as the mean ± SEM. The sample size for in vivo bleomycin fibrosis studies was based on the previous studies in the lab (*6, 84*). No animals were excluded for analysis. Differences in measured variables between experimental and control group were assessed by using Student’s t-tests. One-way followed by Bonferroni’s or two-way ANOVA followed Tukey’s multiple comparison test was used for multiple comparisons. Results were considered statistically significant at P < 0.05, and significance levels were denoted as *P < 0.05, **P < 0.01, ***P < 0.001, and ****P < 0.0001. The number of independent biological replicates is reported as n.

## Acknowledgments

The authors wish to thank the members of the Women’s Guild Lung Institute at Cedars Sinai for their support and helpful discussion during the course of the study. This work was supported by National Institutes of Health grants R01 HL172990 (DJ) and R01 AG078655 (JL and PWN). The UCLA Molecular Screening Shared Resource is supported by Jonsson Comprehensive Cancer Center, award number P30CA016042 by the National Cancer Institute of the National Institutes of Health.

## Author Contributions

DJ and JL conceived and designed the study. AR led the project and performed most of the experiments, analyzed the data, prepared figures, and wrote the paper. YQ and WL performed experiments, analyzed data, prepared figures, and wrote the paper. HH, SY, XZ, XL, NL, GH, PL, and AB took part in mouse, cell culture, and biological experiments. AH, CY, CH, BS, APN, and PWN interpreted data and contributed with comments on the manuscript. TP and PC provided human samples, interpreted data, and contributed with comments on the manuscript. RD helped with drug screening, interpreted data and contributed with comments on the manuscript. AR, YQ, WL, JL, and DJ wrote the paper. All authors read and reviewed the manuscript.

## Materials Availability

Further information and requests for resources and reagents should be directed to and will be fulfilled by the corresponding author, Dianhua Jiang, at, with a completed Materials Transfer Agreement.

---

**Figure S1:**
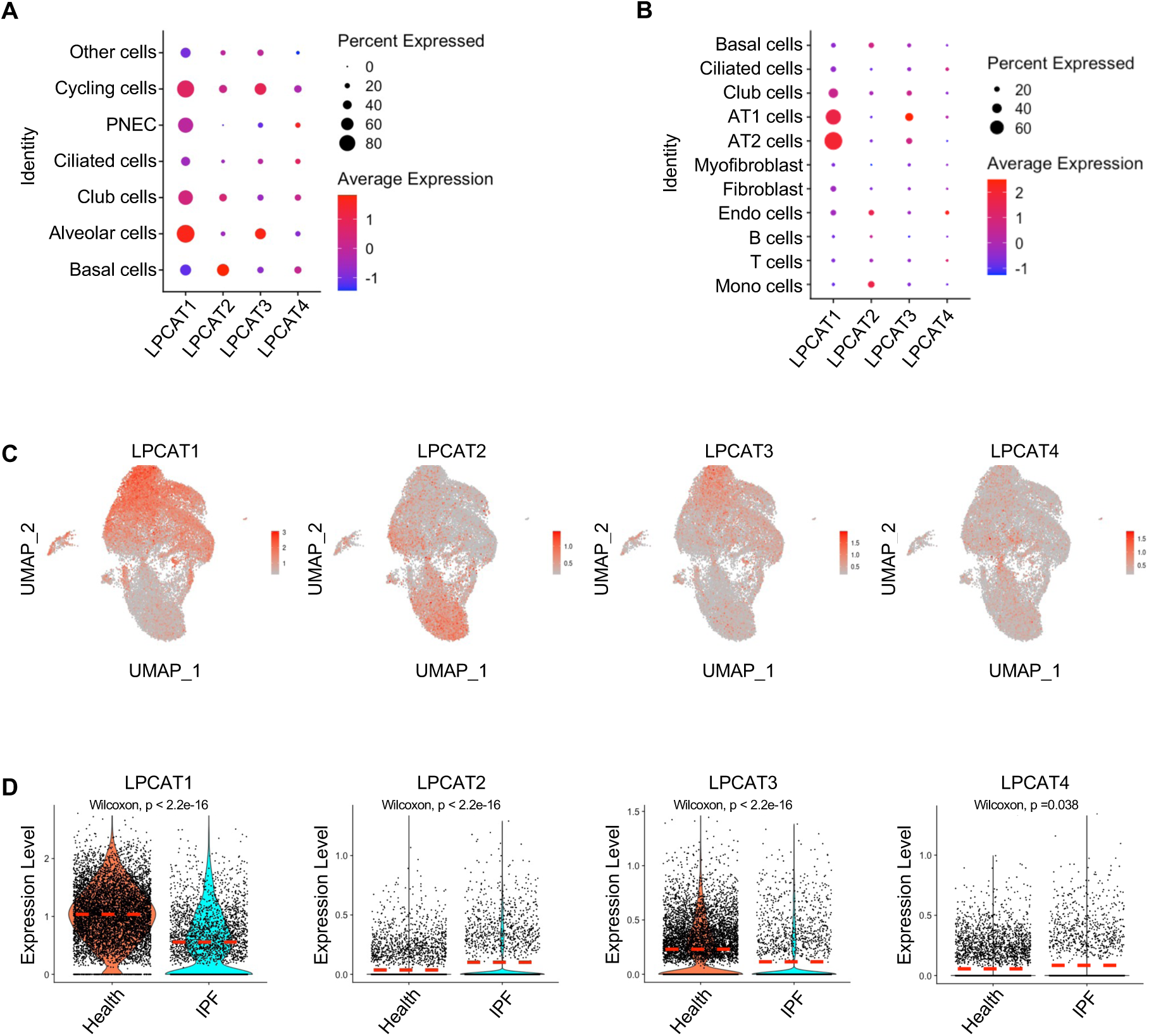
Expression of LPCAT family genes in lung epithelial cells. (A) Dot plot showing the average expression of *LPCAT* family genes in different clusters of epithelial cell types in human lungs, from an in-house scRNA-seq dataset. (B) Dot plot showing the average expression of *LPCAT* family genes in different cell clusters in human lungs from published dataset GSE122960. (C) UMAP visualization of *LPCAT* family gene expression across different epithelial cell clusters from an in-house scRNA-seq dataset. (D) Violin plots showing *LPCAT1* expression levels in AT2s from healthy and IPF lungs from an in-house scRNA-seq dataset.

**Figure S2:**
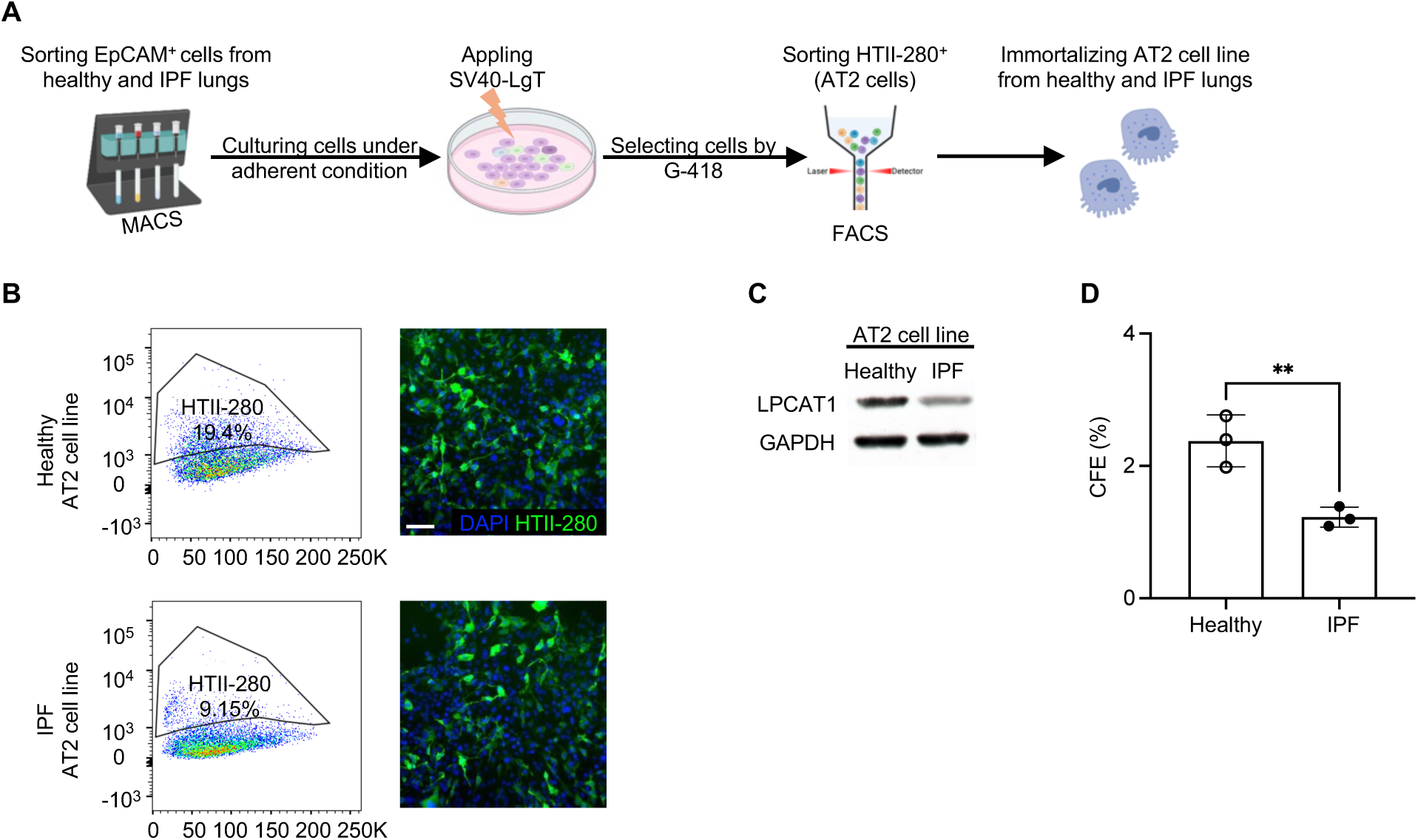
Generation of immortalized AT2 cell lines. (A) Experimental layout for generating immortalized AT2 cell lines from healthy and IPF lungs. (B) Flow-cytometry gating strategies and immunofluorescence staining of the HTII-280+ (green) AT2 marker, sorted from healthy and IPF EpCAM^+^ cells. Scale bars, 100 μm. (C) Representative photographs of Western blot analysis of LPCAT1 and GAPDH in healthy and IPF AT2 cell lines. (D) CFE of immortalized AT2 cell lines from healthy and IPF lungs (n = 3 per group, **p < 0.01).

**Figure S3:**
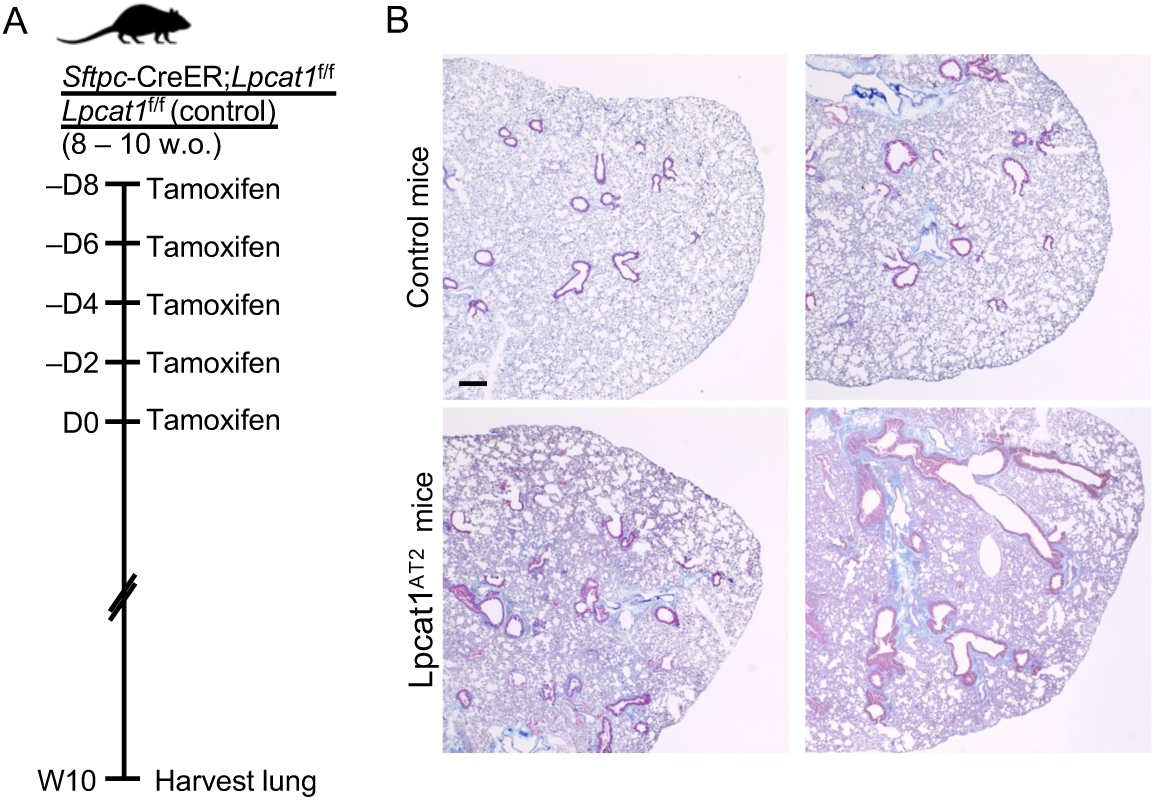
Spontaneous fibrosis in Lpcat1^AT2^ mice. (A) Experimental layout: 8 – 10 weeks old mice were given tamoxifen five times. The lungs were harvested 10 weeks after the last tamoxifen injection. (B) Trichrome staining of lung sections from 18-week-old Lpcat1^AT2^ (10 weeks after the last tamoxifen injection) and control mice. Scale bars: 100 μm. Staining was performed on lung sections from 3 Lpcat1^AT2^ and 3 control mice.

**Figure S4:**
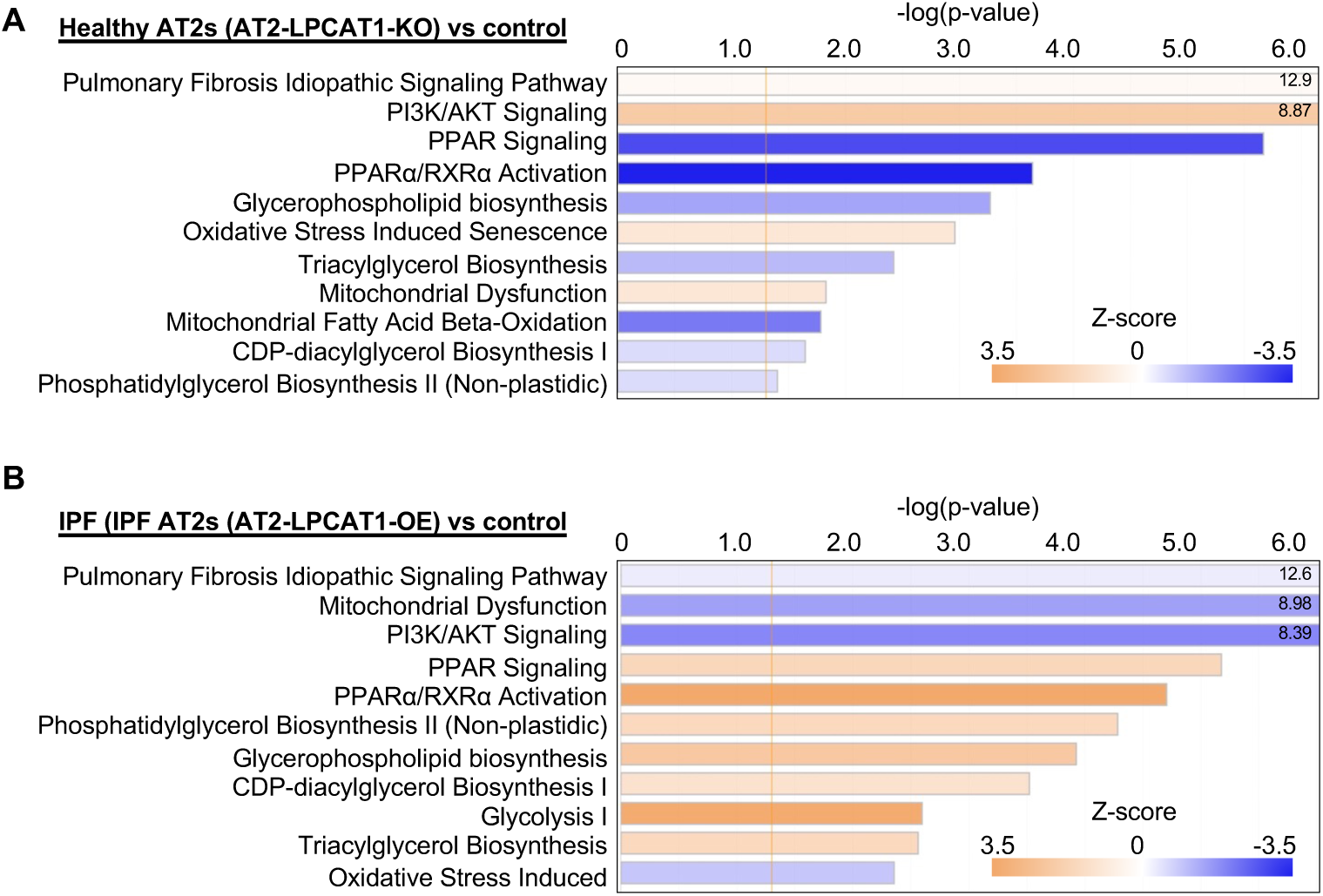
LPCAT1 regulates lipid metabolic pathways. (A) Ingenuity Pathway Analysis (IPA) canonical pathway analysis of healthy AT2-*LPCAT1*-KO cell lines and controls. A negative z-score (blue) indicates a downregulated pathway, whereas a positive z-score (orange) indicates an upregulated pathway. The vertical orange line indicates a Fisher’s exact test p = 0.05 (n = 3 per group). (B) IPA canonical pathway analysis of IPF AT2-*LPCAT1*-OE cell lines and controls. A negative z-score (blue) indicates a downregulated pathway, whereas a positive z-score (orange) indicates an upregulated pathway. The vertical orange line indicates a Fisher’s exact test p = 0.05 (n = 3 per group).

**Figure S5:**
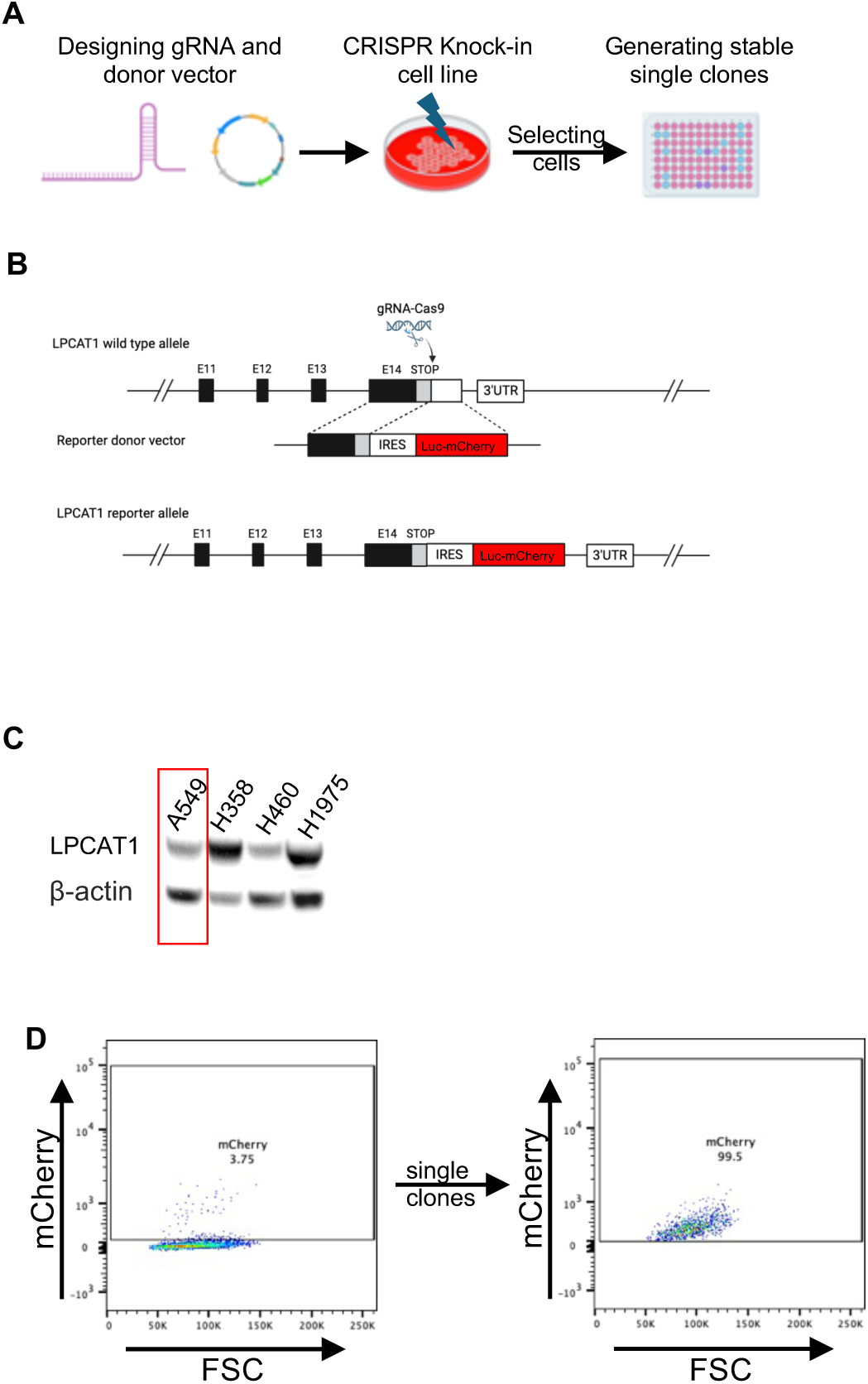
Generation of an LPCAT1 knock-in cell line for drug screening. (A) Experimental layout for generation of *LPCAT1* knock-in cell line. (B) Targeting strategy to generate *LPCAT1* knock-in, tagged with a Luciferase-mCherry fusion allele, using CRISPR/Cas9 system. (C) Western blot analysis of LPCAT1 in different lung cell lines. GAPDH was used as an internal control. (D) Flow-cytometry gating strategies were used to select mCherry+ A549 cells and expand the stable single clones.

